# A chromosome-scale genome and transcriptomic atlas of taro (Colocasia esculenta) corm development

**DOI:** 10.64898/2026.10.11.758265

**Authors:** Xiaolin Cai, Fanglian He

## Abstract

**Aims:** Taro corm development depends on carbohydrate metabolism, but cultivar-level annotation differences complicate the integration of genome comparisons and expression data. We assembled a GY2 reference genome and analysed developmental and tissue transcriptomes to characterise corm-associated transcription in a sequence-aware comparative framework.

**Data resources generated:** The 2.309-Gb GY2 assembly contains 14 pseudomolecules and 20,063 primary gene models. Comparisons with LXY and Bun Long were integrated with six-stage corm and four-tissue RNA-seq datasets. Gene annotations, genomic comparisons, processed expression matrices and analysis scripts are supplied in Data S1, with figure source data in Data S2.

**Key results:** The 14 GY2 pseudomolecules represent 98.83% of the assembly; contig and scaffold N50 values are 11.55 and 182.12 Mb, and liliopsid BUSCO completeness is 97.7%. Genome sequence searches reduced candidate assembly-specific orthogroups from 1,820 to 168. A total of 11,884 genes varied significantly across corm stages. Among 593 strictly corm-enriched genes, 522 were also development-responsive, including nine of 20 selected carbohydrate-associated genes.

**Utility of the resource:** Consistent GY2 v3 gene identifiers link genomic homology, collinearity and gene-level expression, supporting cross-cultivar comparisons and prioritisation of corm-associated candidate genes.

## 1. Introduction

Taro (Colocasia esculenta (L.) Schott) is a vegetatively propagated aroid crop cultivated for its starch-rich corms. Starch accumulation involves sucrose transport and metabolism, plastidial ADP-glucose production, glucan synthesis and remodelling (Braun et al., 2014; Geigenberger, 2011; Seung, 2020; Zeeman et al., 2010). Previous taro transcriptomic and proteomic studies associated corm expansion with starch and sucrose metabolism (Dong et al., 2021; Zou et al., 2025). However, comparative and expression analyses remain difficult without a consistent cultivar-level gene annotation.

The LXY and Bun Long chromosome-scale reference genomes span 2.405 and 2.337 Gb and contain 28,695 and 28,253 annotated genes, respectively (Yin et al., 2021; Sun et al., 2026). Differences in predicted gene content can arise from cultivar variation, repeat resolution and annotation methods; direct genome sequence comparisons are therefore needed to assess apparent gene absence.

Available genomes of Amorphophallus konjac, Cryptocoryne crispatula and Anthurium amnicola extend this comparative context across Araceae (Gao et al., 2022; Qian et al., 2024; Lyu et al., 2025).

Here, we integrate the GY2 genome with three-cultivar sequence comparisons, seven-proteome orthology analysis, a six-stage GY2 corm transcriptome and four-tissue expression profiles. We use 20 annotated carbohydrate-associated genes to examine how genome conservation relates to developmental and tissue-biased transcription.

## 2. Materials and methods

### 2.1 Genome sequencing, assembly and annotation

High-molecular-weight DNA from GY2 was sequenced using PacBio HiFi and Hi-C. HiFi reads were assembled with hifiasm v0.25.0 (Cheng et al., 2021). Hi-C data were processed with Juicer, scaffolded with 3D-DNA and inspected with Juicebox (Dudchenko et al., 2017; Durand et al., 2016a, 2016b). Assembly quality was assessed with BUSCO using the liliopsida dataset, Merqury and the LTR Assembly Index (Manni et al., 2021; Ou et al., 2018; Rhie et al., 2020). Repetitive sequences were annotated with RepeatMasker using rmblastn v2.10.0+ against Dfam 3.1 and RepBase 20181026. Protein-coding annotation integrated transcript evidence from 68 GY2 RNA-seq libraries with same-species protein evidence from the LXY and Bun Long annotations. Protein-to-genome alignments were generated with miniprot (Li, 2023); candidate models were reconciled with transcript and protein evidence, checked for coherent coding structure and validated with gffread. One representative protein and coding sequence was designated for each primary gene.

### 2.2 Taro genome comparison, collinearity and sequence recovery

The GY2, LXY and Bun Long assemblies were compared using minimap2, SyRI v1.7.1, reciprocal MUMmer4/Assemblytics alignments and MCScanX (Goel et al., 2019; Li, 2018; Marçais et al., 2018; Nattestad & Schatz, 2016; Wang et al., 2012). Directional genome alignments required mapping quality ≥20, alignment length ≥1 kb and sequence identity ≥90%; N gaps and rearrangement-boundary masks were excluded. Non-N GY2 sequence aligned reciprocally to both comparators was classified as core, sequence aligned to only one comparator as shared, and sequence without qualifying alignments as candidate-specific. MCScanX identified collinear anchors, and synonymous substitutions per synonymous site (Ks) were estimated by the NG86 method for one-to-one syntenic pairs (Nei & Gojobori, 1986). Miniprot searches were used to locate genomic homologues of orthogroups lacking annotated GY2 genes. Recovery was assessed using stringent and broader sequence-support criteria (Table S2).

### 2.3 Orthology, phylogenomic relationships and gene-family evolution

OrthoFinder was used to infer orthogroups and a seven-proteome tree from GY2, LXY, Bun Long, Anthurium amnicola, Amorphophallus konjac, Cryptocoryne crispatula and Oryza sativa (Emms & Kelly, 2019). For CAFE5 gene-family analysis, the three taro cultivar annotations were represented by their median family count at the Colocasia terminal, with separate analyses using each cultivar count to evaluate sensitivity to annotation differences (Mendes et al., 2020). Family-level P values were adjusted by the Benjamini–Hochberg method. Branch-site codeml tests of filtered single-copy orthogroups designated the Colocasia stem as foreground; candidate groups required an adjusted q < 0.05, at least one Bayes empirical Bayes site with posterior probability ≥0.95 and convergence across optimisation starts (Yang, 2007).

### 2.4 Developmental and tissue RNA-seq resources

The developmental dataset comprised 18 GY2 corm RNA-seq libraries from six stages corresponding to 1, 2, 3, 4, 5 and 8 months after planting (approximately 30, 60, 90, 120, 150 and 240 d, respectively; three biological replicates per stage; PRJNA756190; Dong et al., 2021). The tissue dataset comprised 12 libraries from corm, leaf, petiole and root (three biological replicates per tissue; PRJNA1493089). Reads were aligned to the GY2 reference with STAR in two-pass mode (Dobin et al., 2013) and assigned to 20,063 primary gene models with featureCounts (Liao et al., 2014). Reverse-stranded and unstranded counting were used for the developmental and tissue datasets, respectively, according to read-assignment results. Count matrices were analysed in R 4.5.1 with DESeq2 v1.50.2 (Love et al., 2014), using stage or tissue as the design factor and an intercept-only null model for likelihood-ratio testing. Variance-stabilised values were used for principal-component and expression analyses. After filtering, 17,174 developmental genes and 17,885 tissue genes were tested; 17,142 genes had complete statistics in every adjacent-stage contrast.

### 2.5 Developmental transitions and co-expression analysis

Development-responsive genes were defined by BH-adjusted q < 0.05 in the omnibus likelihood-ratio test. Adjacent-stage differentially expressed genes required BH-adjusted q < 0.05 and |log2 fold change| ≥1. Gene Ontology Biological Process and KEGG over-representation analyses used one-sided Fisher exact tests, with backgrounds restricted to testable genes represented in the corresponding annotation. Terms required at least five background and two foreground genes; BH correction was applied within each transition, direction and database. For GO terms with foreground-gene Jaccard similarity ≥0.70, the term with the lowest adjusted q was retained. WGCNA v1.74 analysed 16,715 genes after exclusion of zero-MAD features (Langfelder & Horvath, 2008). The smallest soft-threshold power meeting scale-free topology R^2^ ≥0.80 and mean connectivity ≥5 was 16. Module eigengenes were correlated with stage order using Spearman correlation. Module membership (kME) was calculated as the Pearson correlation between each eligible gene’s 18-sample VST profile and its assigned module eigengene; BH adjustment was applied across 12,944 non-grey gene tests. High membership required |kME| ≥0.80 and adjusted q < 0.05.

### 2.6 Tissue bias and carbohydrate-pathway integration

Corm expression was compared separately with leaf, petiole and root. Strict corm enrichment required log2 fold change >1 and adjusted q < 0.05 in all three contrasts; a broader corm-biased set required positive corm effects in all three and adjusted q < 0.05 in at least two. Development-responsive genes were intersected with the strict corm-enrichment set. Twenty carbohydrate-associated genes were curated from Swiss-Prot, Gene Ontology and KEGG annotations. Enrichment of these genes in the developmental–corm intersection and WGCNA modules was assessed by Fisher’s exact test. The developmental and tissue heatmaps show independently row-standardised group-mean variance-stabilised expression values.

### 2.7 Three-genome conservation of carbohydrate genes

Orthogroup membership and retained MCScanX anchors were used to classify the 20 carbohydrate-associated genes in LXY and Bun Long. A single in-orthogroup collinear anchor was classified as one-to-one synteny, whereas additional or alternative collinear anchors indicated a duplicated syntenic context. Orthogroup copy counts refer to annotated protein models rather than validated genomic copy number.

## 3. Resource overview

The resource links GY2 v3 gene annotations to sequence-based cultivar comparisons, orthogroup and synteny assignments, and developmental and tissue expression profiles. Data S1 contains processed results and analysis scripts, Data S2 contains source data for Figures 1–4, and Tables S1–S10 present supporting statistics. The raw sequence accessions are specified under Data availability.

**Figure 1.**
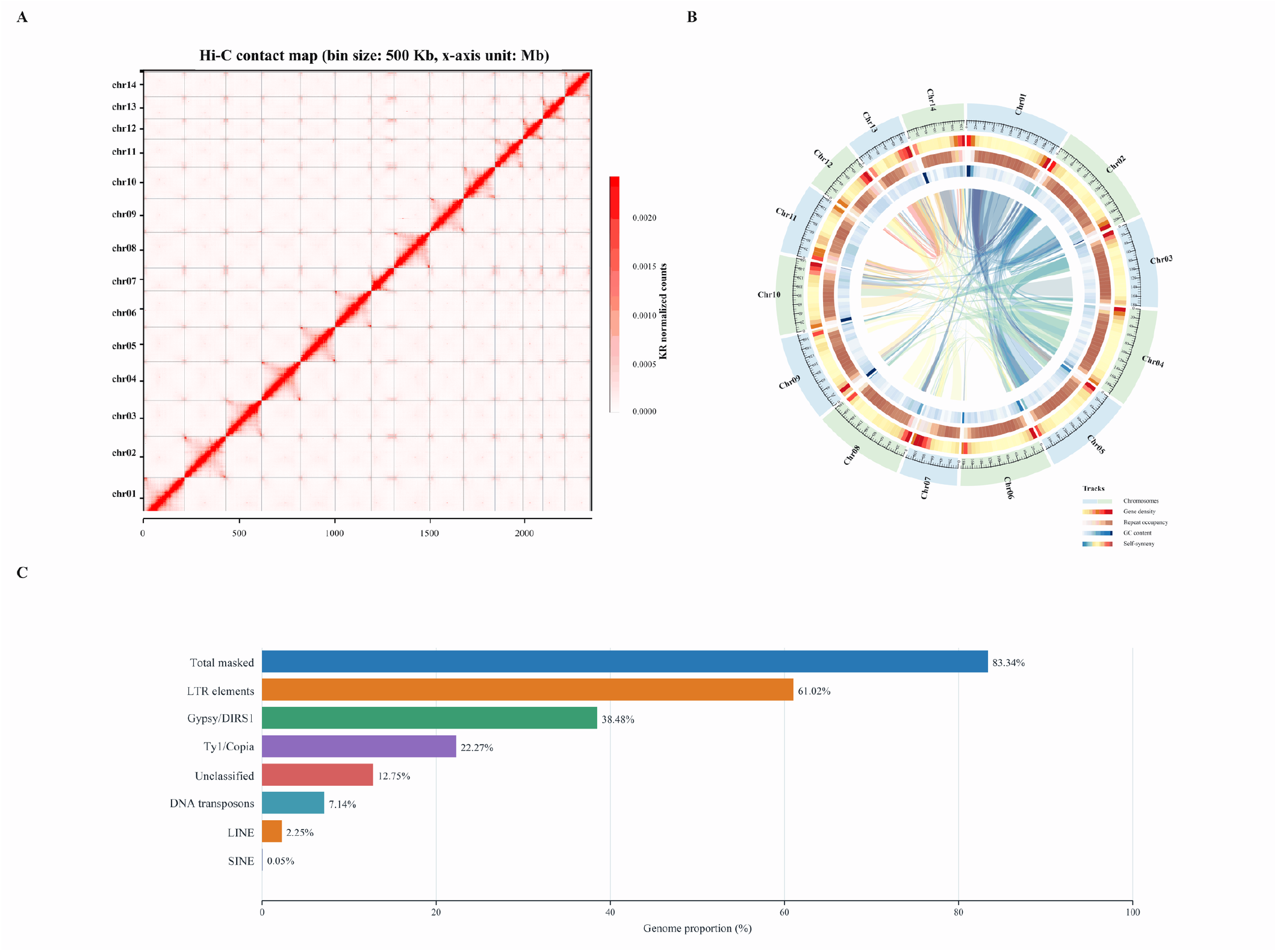
Chromosome-scale assembly and genomic organisation of GY2. (A) Hi-C contact map of the 14 pseudomolecules. (B) Circular genome overview showing chromosomal distributions of gene density, repeat occupancy, GC content and intragenomic syntenic links. (C) Fractions of the GY2 assembly assigned to major repeat categories. LTR subclasses are components of the total LTR fraction and should not be summed with it.

## 4. Results and minor discussion

### 4.1 A chromosome-scale GY2 reference resolves a repeat-rich taro genome

PacBio HiFi sequencing yielded 7,316,334 reads (147.276 Gb; read N50 20,137 bp), and Hi-C sequencing yielded 912,994,097 reads (273.405 Gb; Q30 95.58%). The assembled GY2 genome spans 2,308,659,325 bp, with 98.83% assigned to 14 pseudomolecules (Figure 1A). The contig N50 was 11.55 Mb and the scaffold N50 was 182.12 Mb. Liliopsid BUSCO completeness was 97.7%, the Merqury quality value was 71.62 and the LTR Assembly Index was 12.23 (Table 1).

**Table 1.** Summary of the GY2 genome assembly and annotation.

| Metric | GY2 |
| --- | --- |
| Assembly size (bp) | 2,308,659,325 |
| Primary pseudomolecules | 14 |
| Sequence anchored to 14 pseudomolecules | 98.83% |
| Contig N50 | 11.55 Mb |
| Scaffold N50 | 182.12 Mb |
| Merqury consensus QV | 71.62 |
| LTR Assembly Index | 12.23 |
| Complete BUSCOs (liliopsida) | 97.7% |
| Primary protein-coding genes | 20,063 |

Repetitive sequences accounted for 83.34% of the genome, including 61.02% long terminal repeat retrotransposons. Gypsy/DIRS1 and Ty1/Copia elements contributed 38.48% and 22.27%, respectively (Figure 1C; Table S1). Gene density, repeat abundance and GC content varied across the 14 pseudomolecules, which also contained intragenomic collinear regions (Figure 1B). Annotation identified 20,063 primary protein-coding genes and 23,785 complete transcripts; median gene span, coding-sequence length, exon length and intron length were 4,751, 1,170, 154 and 253 bp, respectively (Table S1).

### 4.2 Direct sequence recovery reduces annotation-driven gene-content discordance

Among 20,484 orthogroups in the three-cultivar comparison, 6,529 lacked annotated GY2 genes. Genome-level searches identified intact coding-sequence support in 4,265 groups, additional homologous sequence in 1,831, partial or fragmented matches in 338, and no detectable match in 95. The stringent and broader sets shared by all three cultivars contained 17,101 and 19,614 orthogroups, respectively. Sequence-based reconciliation reduced candidate assembly-specific orthogroups from 1,820 to 168 (Table S2).

Reciprocal assembly alignments classified 43.67% of GY2 non-N sequence as shared with both comparator assemblies, 41.48% as shared with one and 14.85% as candidate-specific. The corresponding candidate-specific fraction in LXY was 36.27% (Table S2).

Homologous genomic sequence was detected for most orthogroups without an annotated GY2 gene, indicating that annotation-defined absence should not be equated with loss of the underlying sequence. Candidate-specific sequence regions require further assessment across additional accessions.

### 4.3 Taro cultivars retain extensive chromosome-scale conservation

In the seven-proteome OrthoFinder tree, GY2 clustered with Bun Long (node support = 0.942), with LXY sister to this pair (Figure 2A). The distribution of orthogroup-assigned gene categories is shown in Figure 2B. Chromosome comparisons revealed extensive macrosynteny (Figure 2C). Whole-genome alignments covered 75.14% of GY2 against LXY and 88.64% against Bun Long, with callable fractions of 69.94% and 86.70%, respectively. MCScanX identified 20,498 anchors in 676 GY2–LXY blocks and 22,525 anchors in 641 GY2–Bun Long blocks, corresponding to 78.4% and 83.3% of GY2 genes (Table S2).

**Figure 2.**
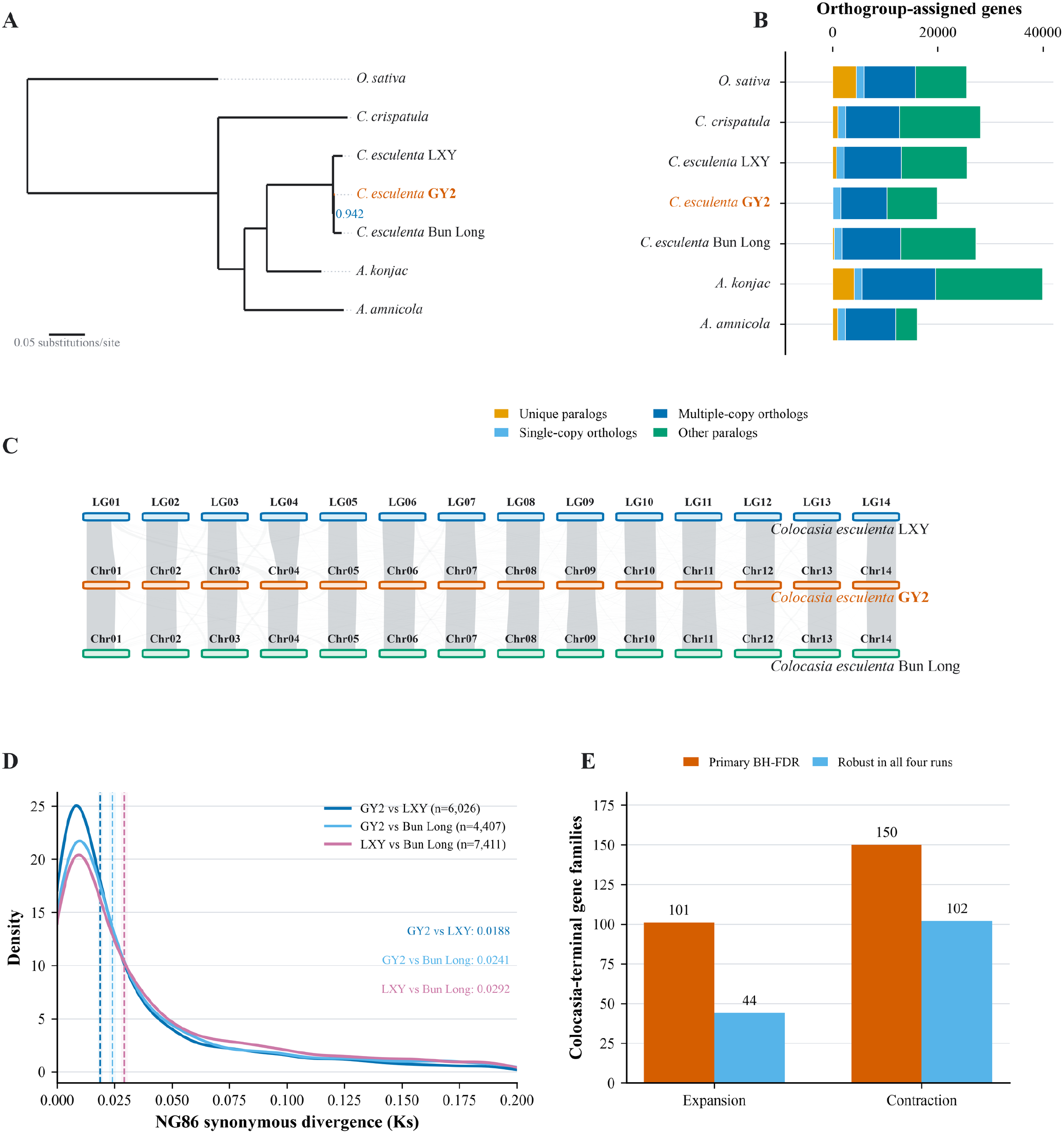
Taro genome relationships and conservation. (A) OrthoFinder tree of seven proteomes (GY2–Bun Long node support = 0.942); branch lengths denote substitutions per site, not elapsed time. (B) Orthogroup-assigned gene categories. (C) MCScanX chromosome collinearity among GY2, LXY and Bun Long. (D) NG86 Ks distributions for one-to-one syntenic pairs (displayed range: 0–0.2); dashed lines mark medians calculated from all finite, unsaturated pairs, and n gives the numbers of those pairs. (E) CAFE5-inferred family expansions and contractions at the Colocasia terminal, showing BH-FDR-significant primary counts and direction-consistent counts across cultivar representations.

Median NG86 Ks values among one-to-one syntenic pairs were 0.0188 for GY2–LXY (n = 6,026), 0.0241 for GY2–Bun Long (n = 4,407) and 0.0292 for LXY–Bun Long (n = 7,411) (Figure 2D; Table S2). Within GY2, 7,720 collinear anchors in 849 blocks showed a median Ks of 0.896, consistent with an older duplication component. In the primary CAFE5 analysis, 101 gene families were inferred to be expanded and 150 contracted at the Colocasia terminal; expansion or contraction direction was retained for 44 and 102 families, respectively, across cultivar-count sensitivity analyses (Figure 2E; Table S3).

### 4.4 Developmental and tissue transcriptomes provide complementary views of corm biology

Principal-component analysis distinguished developmental corm samples (PC1 = 51.7%; PC2 = 15.8%) and the four tissue groups (PC1 = 65.2%; PC2 = 20.1%) (Figure 3A,B). Among 17,174 developmental genes tested by DESeq2, 11,884 differed significantly across stages in the omnibus test (adjusted q < 0.05). Each adjacent-stage contrast included 17,142 genes with complete test statistics (Table S4).

**Figure 3.**
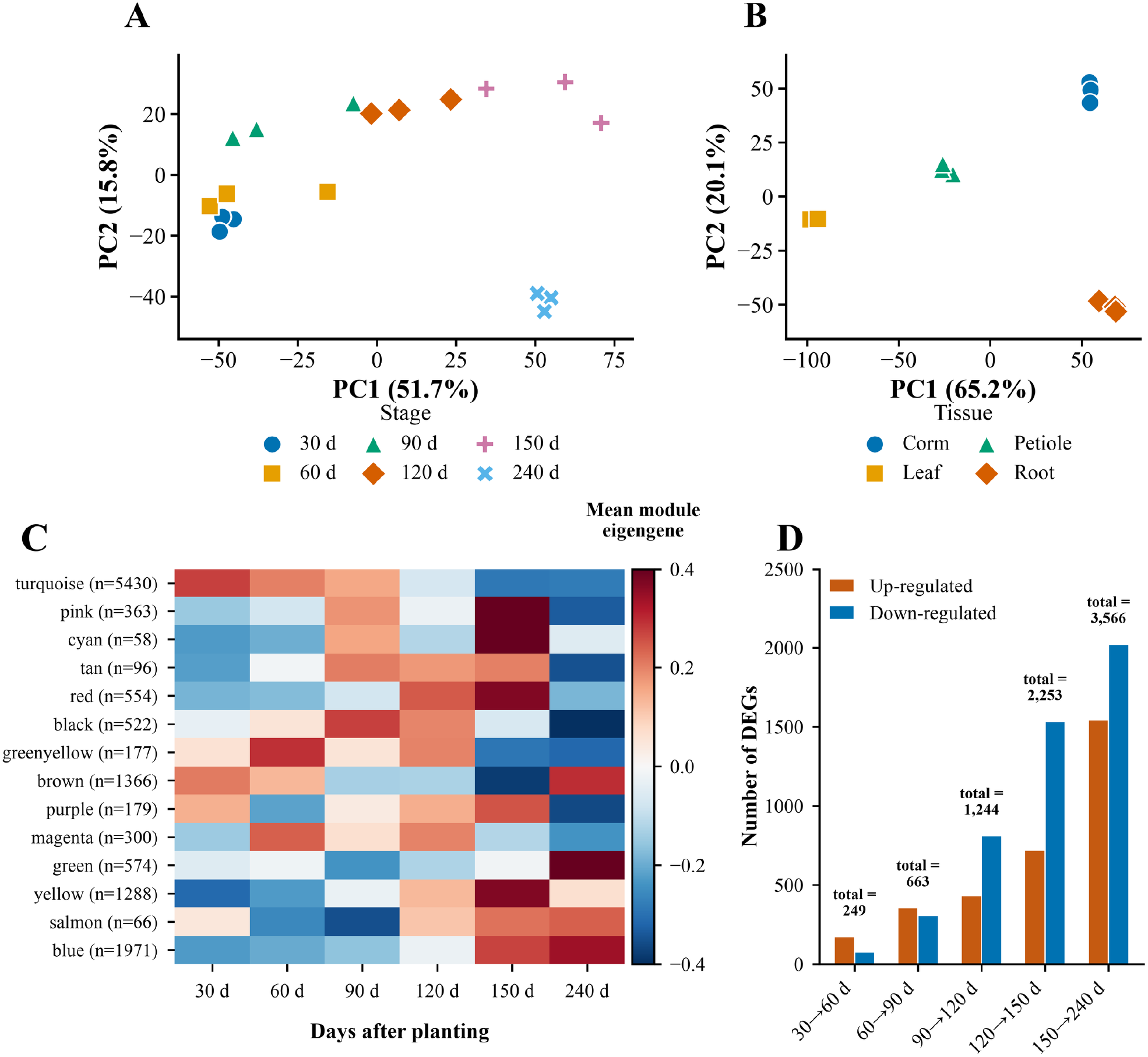
Developmental and tissue transcriptomic profiles in GY2. (A) Principal-component analysis (PCA) of corm samples at six developmental stages. (B) PCA of corm, leaf, petiole and root samples. (C) Stage-mean eigengenes for the 14 non-grey WGCNA modules. (D) Numbers of up- and down-regulated genes in five consecutive developmental contrasts. Three biological replicates were analysed per stage and tissue.

WGCNA analysed 16,715 genes, yielding 14 non-grey modules with distinct developmental eigengene profiles (Figure 3C; Table S5). The five consecutive stage contrasts identified 249, 663, 1,244, 2,253 and 3,566 differentially expressed genes, respectively (Figure 3D; Table S4).

Of 17,885 testable tissue-expressed genes, 593 met the strict criterion for corm enrichment relative to leaf, petiole and root; 522 of these were also development-responsive (Table S6).

### 4.5 Developmental and tissue expression of conserved carbohydrate-associated genes

All 20 GY2 genes selected for sucrose metabolism, ADP-glucose production, starch biosynthesis or glucan remodelling showed significant developmental expression variation, while their expression patterns differed across tissues (Figure 4; Table S7).

**Figure 4.**
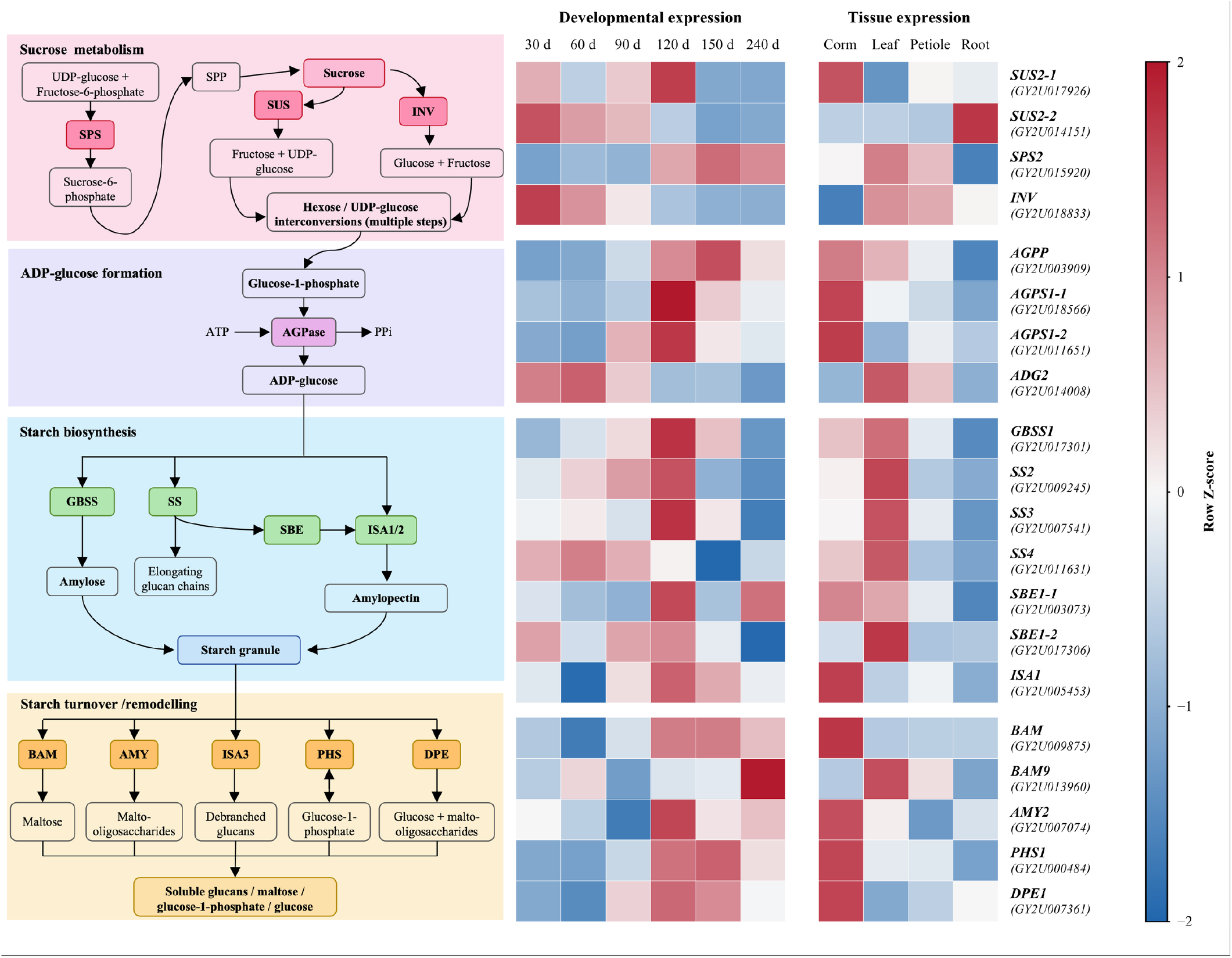
Carbohydrate-associated genes in the GY2 corm transcriptomic atlas. The schematic (left) summarises sucrose metabolism, ADP-glucose production, starch biosynthesis and glucan remodelling. Row-standardised expression values for 20 selected genes are shown across six corm stages (middle) and four tissues (right); standardisation was performed separately for each dataset. Arrows denote annotated pathway relationships, not measured flux.

Nine genes were both development-responsive and strictly corm-enriched: GY2U017926 (SUS2), GY2U018566 and GY2U011651 (AGPS1), GY2U003909 (AGPP), GY2U005453 (ISA1), GY2U009875 (BAM), GY2U007074 (AMY2), GY2U000484 (PHS1) and GY2U007361 (DPE1). Relative to the tested gene background, this intersection was enriched among the 20 curated genes (Fisher’s exact test, odds ratio = 25.89, P = 3.17 × 10□^?^; Table S7). Nineteen genes belonged to non-grey WGCNA modules, with enrichment in the yellow and red modules (Table S8).

All 20 genes had annotated homologues in LXY and Bun Long. Nine had one-to-one syntenic assignments, and eleven belonged to duplicated syntenic contexts (Table S9). These classifications describe conserved genomic context rather than experimentally verified functional equivalence.

## 5. Key conclusions

Genome-level homology recovery demonstrates why apparent gene absence across taro cultivar annotations requires sequence-level assessment. The matched GY2 expression resources identify stage-responsive and corm-enriched carbohydrate-associated candidates for future functional investigation.

## Author contributions

X.C. and F.H. conceived the study. F.H. coordinated the biological work, supervised the project and acquired funding. X.C. curated the genomic and transcriptomic resources, performed the comparative and integrative analyses and visualisation, and drafted the manuscript. Both authors interpreted the results, revised the manuscript and approved the final version.

## Acknowledgements

We thank the staff of the Guangxi Academy of Agricultural Sciences for plant maintenance and sample collection and the sequencing service teams for generation of genomic and RNA-sequencing data.

## Data availability and FAIR statement

The GY2 genome sequence was submitted to NCBI under BioProject PRJNA1492760, BioSample SAMN61520667 and WGS accession JCAWEZ000000000. The developmental and tissue RNA-seq datasets are associated with PRJNA756190 and PRJNA1493089, respectively. Data S1 contains annotated gene models, processed expression matrices, comparative-genomic results and analysis scripts; Data S2 provides source data for Figures 1–4, and Tables S1–S10 contain supporting statistics. Tab-delimited files use the GY2 v3 gene identifiers to enable integration across datasets.

## Generative AI statement

OpenAI ChatGPT was used to assist with manuscript language editing and the development of analysis and plotting scripts. The authors reviewed the resulting text and code, verified the reported results, and remain responsible for the scientific content.

## Supporting information

Table S1. Sequencing input, repeat annotation and gene-model structural details for GY2.

Table S2. Cultivar comparative genomics, assembly-to-assembly differences and sequence-supported orthogroup recovery.

Table S3. Araceae orthology, Colocasia gene-family evolution and high-confidence branch-site candidates. Table S4. Adjacent-stage differential-expression summary for the six-stage corm series.

Table S5. Non-grey WGCNA module sizes, stage associations and eigengene means. Table S6. Strictly corm-enriched genes with developmental-response status.

Table S7. Integrated developmental, tissue, network and functional-annotation evidence for 20 curated starch/sucrose genes.

Table S8. WGCNA module enrichment of the curated starch/sucrose gene set.

Table S9. Three-genome orthogroup and synteny classification for the 20 curated starch/sucrose genes.

Table S10. Non-redundant GO Biological Process and KEGG enrichment across adjacent developmental transitions.

Data S1. GY2 gene annotation, processed genomic and transcriptomic results, and analysis scripts. Data S2. Source data for Figures 1–4.

