## Supplementary material for "A chromosome-scale genome and transcriptomic atlas of taro (Colocasia esculenta) corm development": Data S1: README.md

# Data S1. GY2 genome and transcriptome resources

**Reference annotation:** GY2 v3, 20,063 primary genes.  
**Developmental RNA-seq:** PRJNA756190, six stages, three biological replicates per stage.  
**Tissue RNA-seq:** PRJNA1493089, four tissues, three biological replicates per tissue.

| Directory | Contents |
| --- | --- |
| `inputs/annotation/` | Primary GY2 gene annotation, protein sequences and gene ID mapping |
| `inputs/expression/` | Gene counts, sample metadata, DESeq2 results and variance-stabilised expression |
| `inputs/developmental/results/` | Stage contrasts, WGCNA modules, PCA scores and expression matrices |
| `inputs/functional_annotation/` | Gene Ontology and KEGG assignments |
| `inputs/comparative/` | Orthogroups and collinear anchors |
| `inputs/pathway/` | Carbohydrate-associated gene selection and annotation |
| `results/tables/` | Differential-expression, co-expression, enrichment and gene-conservation tables |
| `scripts/` | R and Python analysis source code |

The gene annotation and all expression matrices use the GY2 v3 gene identifiers. The genome sequence and sequencing reads are referenced by accession numbers in the manuscript's Data availability statement.

## Reproducing the included analyses

The downstream integrated analysis can be rerun from the archive root using `python3 scripts/integrated_expression_analysis.py`. The developmental DESeq2/WGCNA analysis script can be invoked with `Rscript scripts/developmental_expression.R --counts inputs/expression/counts/PRJNA756190.gene_counts.tsv --sample-map inputs/expression/developmental_sample_map.tsv --output-dir output_developmental --annotation-label GY2.uniform_reannotated.v3`. The R analysis requires DESeq2, WGCNA and jsonlite.

The archive contains gene-level counts and downstream analysis inputs, not original FASTQ reads or all assembly and gene-model prediction programs. The tissue sample metadata includes original workspace-specific FASTQ paths as provenance; those paths are not portable. Retrieve corresponding raw data using the accessions in the manuscript once those records are accessible.
