## Supplementary material for "A chromosome-scale genome and transcriptomic atlas of taro (Colocasia esculenta) corm development": Data S1: developmental_expression.R

```
#!/usr/bin/env Rscript

suppressPackageStartupMessages({
  library(DESeq2)
  library(WGCNA)
  library(jsonlite)
})

options(stringsAsFactors = FALSE)
allowWGCNAThreads(nThreads = as.integer(Sys.getenv("WGCNA_THREADS", "32")))

args <- commandArgs(trailingOnly = TRUE)
value_after <- function(flag) {
  position <- match(flag, args)
  if (is.na(position) || position == length(args)) stop("missing argument: ", flag)
  args[[position + 1L]]
}

optional_value_after <- function(flag, default) {
  position <- match(flag, args)
  if (is.na(position)) return(default)
  if (position == length(args)) stop("missing argument: ", flag)
  args[[position + 1L]]
}

counts_path <- value_after("--counts")
sample_map_path <- value_after("--sample-map")
output_dir <- value_after("--output-dir")
annotation_label <- optional_value_after("--annotation-label", "GY2.uniform_reannotated.v3")
dir.create(output_dir, recursive = TRUE, showWarnings = FALSE)
dir.create(file.path(output_dir, "contrasts"), recursive = TRUE, showWarnings = FALSE)

write_tsv <- function(frame, path) {
  write.table(frame, path, sep = "\t", quote = FALSE, row.names = FALSE, na = "NA")
}

bh_safe <- function(values) {
  adjusted <- rep(NA_real_, length(values))
  valid <- is.finite(values)
  adjusted[valid] <- p.adjust(values[valid], method = "BH")
  adjusted
}

counts_frame <- read.delim(counts_path, check.names = FALSE)
sample_map <- read.delim(sample_map_path, check.names = FALSE)
required_map <- c("run", "sample_id", "stage", "replicate")
if (!all(required_map %in% names(sample_map))) stop("sample map lacks required columns")
if (nrow(sample_map) != 18L || anyDuplicated(sample_map$sample_id) || anyDuplicated(sample_map$run)) {
  stop("sample map must contain 18 unique runs and sample IDs")
}
stage_counts <- table(sample_map$stage)
if (!identical(as.integer(stage_counts[paste0("S", 1:6)]), rep(3L, 6L))) {
  stop("sample map does not close to six stages by three replicates")
}
if (!"gene_id" %in% names(counts_frame) || anyDuplicated(counts_frame$gene_id)) {
  stop("count matrix must have unique gene_id rows")
}
if (!all(sample_map$sample_id %in% names(counts_frame))) stop("count matrix lacks mapped samples")

count_matrix <- as.matrix(counts_frame[, sample_map$sample_id, drop = FALSE])
storage.mode(count_matrix) <- "integer"
rownames(count_matrix) <- counts_frame$gene_id
if (any(count_matrix < 0L) || any(!is.finite(count_matrix))) stop("invalid count matrix")

sample_map$stage <- factor(sample_map$stage, levels = paste0("S", 1:6))
rownames(sample_map) <- sample_map$sample_id
eligible <- rowSums(count_matrix >= 10L) >= 3L
filter_audit <- data.frame(
  gene_id = rownames(count_matrix),
  total_count = rowSums(count_matrix),
  samples_count_ge_10 = rowSums(count_matrix >= 10L),
  DESeq2_eligible = eligible,
  stringsAsFactors = FALSE
)
write_tsv(filter_audit, file.path(output_dir, "feature_filter.tsv"))
if (sum(eligible) < 1000L) stop("fewer than 1,000 genes pass the frozen count filter")

dds_base <- DESeqDataSetFromMatrix(
  countData = count_matrix[eligible, , drop = FALSE],
  colData = sample_map,
  design = ~ stage
)
dds_lrt <- DESeq(dds_base, test = "LRT", reduced = ~ 1, quiet = TRUE)
lrt <- as.data.frame(results(dds_lrt, independentFiltering = FALSE))
lrt$gene_id <- rownames(lrt)
lrt <- lrt[, c("gene_id", setdiff(names(lrt), "gene_id"))]
names(lrt)[names(lrt) == "padj"] <- "padj_BH_all_eligible"
lrt$dynamic_BH05 <- is.finite(lrt$padj_BH_all_eligible) & lrt$padj_BH_all_eligible < 0.05
write_tsv(lrt, file.path(output_dir, "omnibus_LRT_all_genes.tsv"))

dds_wald <- DESeq(dds_base, test = "Wald", quiet = TRUE)
contrast_summary <- list()
summary_index <- 1L
for (later_index in 2:6) {
  for (earlier_index in 1:(later_index - 1L)) {
    later <- paste0("S", later_index)
    earlier <- paste0("S", earlier_index)
    label <- paste0(later, "_vs_", earlier)
    result <- as.data.frame(results(
      dds_wald,
      contrast = c("stage", later, earlier),
      independentFiltering = FALSE,
      alpha = 0.05
    ))
    result$gene_id <- rownames(result)
    result <- result[, c("gene_id", setdiff(names(result), "gene_id"))]
    names(result)[names(result) == "padj"] <- "padj_BH_all_eligible"
    result$DE <- (
      is.finite(result$padj_BH_all_eligible)
      & result$padj_BH_all_eligible < 0.05
      & abs(result$log2FoldChange) >= 1
    )
    write_tsv(result, file.path(output_dir, "contrasts", paste0(label, ".all_genes.tsv")))
    write_tsv(result[result$DE, , drop = FALSE], file.path(output_dir, "contrasts", paste0(label, ".DE.tsv")))
    contrast_summary[[summary_index]] <- data.frame(
      contrast = label,
      tested_genes = nrow(result),
      DE_genes = sum(result$DE),
      up_later = sum(result$DE & result$log2FoldChange > 0),
      down_later = sum(result$DE & result$log2FoldChange < 0)
    )
    summary_index <- summary_index + 1L
  }
}
write_tsv(do.call(rbind, contrast_summary), file.path(output_dir, "contrast_summary.tsv"))

normalized <- counts(dds_wald, normalized = TRUE)
normalized_frame <- data.frame(gene_id = rownames(normalized), normalized, check.names = FALSE)
write_tsv(normalized_frame, gzfile(file.path(output_dir, "normalized_counts_deseq2.tsv.gz"), "wt"))

vst_object <- vst(dds_wald, blind = FALSE)
vst_matrix <- assay(vst_object)
vst_frame <- data.frame(gene_id = rownames(vst_matrix), vst_matrix, check.names = FALSE)
write_tsv(vst_frame, gzfile(file.path(output_dir, "vst_expression.tsv.gz"), "wt"))

pca <- prcomp(t(vst_matrix), center = TRUE, scale. = FALSE)
pca_scores <- data.frame(
  sample_id = rownames(pca$x),
  stage = sample_map[rownames(pca$x), "stage"],
  replicate = sample_map[rownames(pca$x), "replicate"],
  pca$x[, seq_len(min(6L, ncol(pca$x))), drop = FALSE],
  check.names = FALSE
)
write_tsv(pca_scores, file.path(output_dir, "PCA_scores.tsv"))
pca_variance <- data.frame(
  component = paste0("PC", seq_along(pca$sdev)),
  variance_fraction = pca$sdev^2 / sum(pca$sdev^2)
)
write_tsv(pca_variance, file.path(output_dir, "PCA_variance.tsv"))
write_tsv(data.frame(sample_id = names(sizeFactors(dds_wald)), size_factor = sizeFactors(dds_wald)),
          file.path(output_dir, "size_factors.tsv"))

mads <- apply(vst_matrix, 1L, mad)
network_ids <- names(mads)[is.finite(mads) & mads > 0]
network_order <- order(-mads[network_ids], network_ids)
network_ids <- network_ids[network_order]
if (length(network_ids) > 20000L) network_ids <- network_ids[seq_len(20000L)]
if (length(network_ids) < 1000L) stop("fewer than 1,000 genes are WGCNA eligible")
dat_expr <- t(vst_matrix[network_ids, , drop = FALSE])
good <- goodSamplesGenes(dat_expr, verbose = 0)
if (!good$allOK) {
  dat_expr <- dat_expr[good$goodSamples, good$goodGenes, drop = FALSE]
}
if (nrow(dat_expr) != 18L) stop("WGCNA sample QC removed a developmental sample")

power_ids <- colnames(dat_expr)[seq_len(min(5000L, ncol(dat_expr)))]
powers <- 1:20
soft <- pickSoftThreshold(
  dat_expr[, power_ids, drop = FALSE],
  powerVector = powers,
  networkType = "signed hybrid",
  corFnc = "bicor",
  corOptions = list(use = "p", maxPOutliers = 0.1),
  verbose = 0
)
fit <- soft$fitIndices
passing_power <- fit$Power[is.finite(fit$SFT.R.sq) & fit$SFT.R.sq >= 0.80 & fit$mean.k. >= 5]
if (length(passing_power)) {
  selected_power <- min(passing_power)
  power_rule <- "smallest_R2_ge_0.80_mean_connectivity_ge_5"
} else {
  fallback <- fit[is.finite(fit$SFT.R.sq) & fit$mean.k. >= 1, , drop = FALSE]
  if (!nrow(fallback)) stop("no soft power has finite fit and mean connectivity >=1")
  selected_power <- fallback$Power[order(-fallback$SFT.R.sq, fallback$Power)][1]
  power_rule <- "fallback_max_R2_among_mean_connectivity_ge_1"
}
fit$selected <- fit$Power == selected_power
write_tsv(fit, file.path(output_dir, "WGCNA_soft_threshold.tsv"))

network <- blockwiseModules(
  dat_expr,
  power = selected_power,
  networkType = "signed hybrid",
  TOMType = "signed",
  corType = "bicor",
  maxPOutliers = 0.1,
  minModuleSize = 30,
  deepSplit = 2,
  mergeCutHeight = 0.25,
  pamRespectsDendro = FALSE,
  maxBlockSize = 20000,
  numericLabels = FALSE,
  saveTOMs = FALSE,
  verbose = 2
)
module_colors <- network$colors
names(module_colors) <- colnames(dat_expr)
module_eigengenes <- orderMEs(moduleEigengenes(dat_expr, colors = module_colors)$eigengenes)
kme <- signedKME(dat_expr, module_eigengenes, corFnc = "bicor", corOptions = "use = 'p'")
own_kme <- vapply(seq_along(module_colors), function(index) {
  column <- paste0("kME", module_colors[[index]])
  if (column %in% colnames(kme)) kme[index, column] else NA_real_
}, numeric(1))
assignments <- data.frame(
  gene_id = names(module_colors),
  module_id = unname(module_colors),
  MAD = mads[names(module_colors)],
  own_module_kME = own_kme,
  stringsAsFactors = FALSE
)
assignments <- assignments[order(assignments$module_id, -assignments$own_module_kME, assignments$gene_id), ]
write_tsv(assignments, file.path(output_dir, "WGCNA_module_assignments.tsv"))

eigengene_frame <- data.frame(sample_id = rownames(module_eigengenes), module_eigengenes, check.names = FALSE)
write_tsv(eigengene_frame, file.path(output_dir, "WGCNA_module_eigengenes.tsv"))

module_rows <- list()
module_names <- colnames(module_eigengenes)
for (index in seq_along(module_names)) {
  module_name <- module_names[[index]]
  values <- module_eigengenes[, module_name]
  stage_model <- anova(lm(values ~ sample_map[rownames(module_eigengenes), "stage"]))
  stage_means <- tapply(values, sample_map[rownames(module_eigengenes), "stage"], mean)
  rho <- suppressWarnings(cor(seq_along(stage_means), as.numeric(stage_means), method = "spearman"))
  module_rows[[index]] <- data.frame(
    module_id = sub("^ME", "", module_name),
    genes = sum(module_colors == sub("^ME", "", module_name)),
    stage_ANOVA_F = stage_model$`F value`[1],
    stage_ANOVA_p = stage_model$`Pr(>F)`[1],
    stage_mean_Spearman_rho = rho,
    peak_stage = names(stage_means)[which.max(stage_means)],
    S1_mean = stage_means[["S1"]], S2_mean = stage_means[["S2"]],
    S3_mean = stage_means[["S3"]], S4_mean = stage_means[["S4"]],
    S5_mean = stage_means[["S5"]], S6_mean = stage_means[["S6"]]
  )
}
module_summary <- do.call(rbind, module_rows)
module_summary$stage_ANOVA_q_BH_modules <- bh_safe(module_summary$stage_ANOVA_p)
write_tsv(module_summary, file.path(output_dir, "WGCNA_module_stage_summary.tsv"))

saveRDS(
  list(dds_lrt = dds_lrt, dds_wald = dds_wald, vst = vst_object,
       network = network, selected_power = selected_power),
  file.path(output_dir, "reannotated_developmental_frozen_objects.rds")
)
capture.output(sessionInfo(), file = file.path(output_dir, "sessionInfo.txt"))

summary <- list(
  status = "APPROVED",
  annotation = annotation_label,
  samples = ncol(count_matrix),
  stages = 6L,
  biological_replicates_per_stage = 3L,
  total_reconstructed_genes = nrow(count_matrix),
  DESeq2_eligible_genes = sum(eligible),
  omnibus_dynamic_BH05 = sum(lrt$dynamic_BH05, na.rm = TRUE),
  pairwise_contrasts = 15L,
  WGCNA_genes = ncol(dat_expr),
  WGCNA_modules_excluding_grey = length(setdiff(unique(module_colors), "grey")),
  WGCNA_grey_genes = sum(module_colors == "grey"),
  soft_power = as.integer(selected_power),
  soft_power_rule = power_rule,
  interpretation = "Developmental coexpression and stage association support prioritization only; regulatory direction and causality are not established."
)
writeLines(toJSON(summary, pretty = TRUE, auto_unbox = TRUE), file.path(output_dir, "developmental_summary.json"))
```
