## Supplementary material for "A chromosome-scale genome and transcriptomic atlas of taro (Colocasia esculenta) corm development": Data S1: integrated_expression_analysis.py

```
#!/usr/bin/env python3
"""Integrate GY2 gene annotation, expression and comparative genomics."""

from __future__ import annotations

import hashlib
import json
import math
import platform
import re
import sys
from collections import Counter, defaultdict
from pathlib import Path

import matplotlib
matplotlib.use("Agg")
import matplotlib.pyplot as plt
from matplotlib.colors import LinearSegmentedColormap, ListedColormap
import numpy as np
import pandas as pd
from scipy.stats import fisher_exact, pearsonr


ROOT = Path(__file__).resolve().parents[1]
INPUT = ROOT / "inputs"
TABLES = ROOT / "results" / "tables"
FIGURES = ROOT / "results" / "figures"
QC = ROOT / "qc"

FDR = 0.05
LFC = 1.0
STAGES = ["S1", "S2", "S3", "S4", "S5", "S6"]
STAGE_DAYS = {"S1": 30, "S2": 60, "S3": 90, "S4": 120, "S5": 150, "S6": 240}
TRANSITIONS = [
    ("S2_vs_S1", "S1", "S2"),
    ("S3_vs_S2", "S2", "S3"),
    ("S4_vs_S3", "S3", "S4"),
    ("S5_vs_S4", "S4", "S5"),
    ("S6_vs_S5", "S5", "S6"),
]

BLUE = "#0072B2"
ORANGE = "#D55E00"
GREEN = "#009E73"
SKY = "#56B4E9"
PURPLE = "#CC79A7"
YELLOW = "#E69F00"
INK = "#222222"
MUTED = "#6B7280"
LIGHT = "#E5E7EB"


def ensure_dirs() -> None:
    for path in (TABLES, FIGURES, QC):
        path.mkdir(parents=True, exist_ok=True)


def read_tsv(path: Path, **kwargs) -> pd.DataFrame:
    if not path.exists():
        raise FileNotFoundError(path)
    return pd.read_csv(path, sep="\t", low_memory=False, **kwargs)


def write_tsv(df: pd.DataFrame, path: Path) -> None:
    df.to_csv(path, sep="\t", index=False, na_rep="")


def bh_adjust(values) -> np.ndarray:
    p = np.asarray(values, dtype=float)
    out = np.full(p.shape, np.nan, dtype=float)
    ok = np.isfinite(p)
    if not ok.any():
        return out
    vals = p[ok]
    order = np.argsort(vals)
    ranked = vals[order]
    n = len(ranked)
    q = ranked * n / np.arange(1, n + 1)
    q = np.minimum.accumulate(q[::-1])[::-1]
    q = np.clip(q, 0, 1)
    restored = np.empty(n, dtype=float)
    restored[order] = q
    out[np.where(ok)[0]] = restored
    return out


def sha256(path: Path) -> str:
    h = hashlib.sha256()
    with path.open("rb") as handle:
        for chunk in iter(lambda: handle.read(1024 * 1024), b""):
            h.update(chunk)
    return h.hexdigest()


def strip_tx(value: str) -> str:
    return re.sub(r"\.t\d+$", "", str(value).strip())


def split_members(value) -> list[str]:
    if pd.isna(value) or not str(value).strip():
        return []
    return [x.strip() for x in str(value).split(",") if x.strip()]


def odds_ci(a: int, b: int, c: int, d: int) -> tuple[float, float, float]:
    # Haldane-Anscombe correction produces a finite descriptive CI.
    aa, bb, cc, dd = [x + 0.5 for x in (a, b, c, d)]
    estimate = (aa * dd) / (bb * cc)
    se = math.sqrt(1 / aa + 1 / bb + 1 / cc + 1 / dd)
    lo = math.exp(math.log(estimate) - 1.96 * se)
    hi = math.exp(math.log(estimate) + 1.96 * se)
    return estimate, lo, hi


def save_figure(fig: plt.Figure, stem: str) -> None:
    fig.savefig(FIGURES / f"{stem}.tiff", dpi=600, bbox_inches="tight",
                pil_kwargs={"compression": "tiff_lzw"})
    fig.savefig(FIGURES / f"{stem}_preview.png", dpi=180, bbox_inches="tight")
    plt.close(fig)


def parse_go(path: Path, author_to_v3: dict[str, str]) -> tuple[dict[str, set], dict[str, dict]]:
    term_genes: dict[str, set] = defaultdict(set)
    metadata: dict[str, dict] = {}
    with path.open(errors="replace") as handle:
        for line in handle:
            if line.startswith("#") or not line.strip():
                continue
            parts = line.rstrip("\n\r").split("\t")
            old_id = parts[0]
            gene = author_to_v3.get(old_id)
            if not gene:
                continue
            for annotation in parts[2:]:
                match = re.search(r"^(.*?):\s*(.*?)\s*\((GO:\d+)\)", annotation.strip())
                if not match:
                    continue
                ontology, name, term_id = match.groups()
                term_genes[term_id].add(gene)
                metadata[term_id] = {"ontology": ontology, "term_name": name}
    return term_genes, metadata


def parse_kegg(path: Path, author_to_v3: dict[str, str]) -> tuple[dict[str, set], dict[str, dict]]:
    term_genes: dict[str, set] = defaultdict(set)
    metadata: dict[str, dict] = {}
    with path.open(errors="replace") as handle:
        header = next(handle, None)
        for line in handle:
            parts = line.rstrip("\n\r").split("\t")
            if len(parts) < 4:
                continue
            name, term_id, _, proteins = parts[:4]
            for old_id in proteins.split(";"):
                gene = author_to_v3.get(old_id.strip())
                if gene:
                    term_genes[term_id].add(gene)
            metadata[term_id] = {"ontology": "KEGG pathway", "term_name": name}
    return term_genes, metadata


def enrich_one(foreground: set[str], background: set[str], term_genes: dict[str, set],
               metadata: dict[str, dict], transition: str, direction: str,
               database: str) -> pd.DataFrame:
    rows = []
    annotated_genes = set().union(*term_genes.values()) if term_genes else set()
    analysis_background = background & annotated_genes
    n = len(analysis_background)
    fg = foreground & analysis_background
    for term_id, members in term_genes.items():
        in_bg = members & analysis_background
        a = len(in_bg & fg)
        k = len(in_bg)
        if k < 5 or a < 2:
            continue
        b = len(fg) - a
        c = k - a
        d = n - a - b - c
        if min(a, b, c, d) < 0:
            continue
        odds, p = fisher_exact([[a, b], [c, d]], alternative="greater")
        fold = (a / len(fg)) / (k / n) if fg and k else np.nan
        rows.append({
            "database": database,
            "transition": transition,
            "direction": direction,
            "ontology": metadata[term_id]["ontology"],
            "term_id": term_id,
            "term_name": metadata[term_id]["term_name"],
            "n_background": n,
            "n_foreground": len(fg),
            "background_hits": k,
            "foreground_hits": a,
            "fold_enrichment": fold,
            "odds_ratio": odds,
            "p_value": p,
            "foreground_gene_ids": ";".join(sorted(in_bg & fg)),
        })
    result = pd.DataFrame(rows)
    if not result.empty:
        result["q_value"] = bh_adjust(result["p_value"])
        result = result.sort_values(["q_value", "p_value", "term_id"]).reset_index(drop=True)
    return result


def reduce_redundancy(go: pd.DataFrame, kegg: pd.DataFrame) -> pd.DataFrame:
    retained = []
    sig_go = go[go["q_value"] < FDR].copy()
    for _, group in sig_go.groupby(["transition", "direction", "ontology"], sort=False):
        kept_sets: list[set[str]] = []
        for _, row in group.sort_values(["q_value", "p_value"]).iterrows():
            genes = set(str(row["foreground_gene_ids"]).split(";"))
            redundant = False
            for prior in kept_sets:
                union = genes | prior
                jaccard = len(genes & prior) / len(union) if union else 0
                if jaccard >= 0.70:
                    redundant = True
                    break
            if not redundant:
                record = row.to_dict()
                record["representative_rule"] = "lowest q retained; Jaccard gene overlap < 0.70"
                retained.append(record)
                kept_sets.append(genes)
            if len(kept_sets) >= 5:
                break
    sig_kegg = kegg[kegg["q_value"] < FDR].copy()
    for _, group in sig_kegg.groupby(["transition", "direction"], sort=False):
        for _, row in group.sort_values(["q_value", "p_value"]).head(5).iterrows():
            record = row.to_dict()
            record["representative_rule"] = "top five significant KEGG pathways per transition/direction"
            retained.append(record)
    return pd.DataFrame(retained)


def parse_gff_coordinates(path: Path) -> dict[str, tuple[str, int, int, str]]:
    coords: dict[str, tuple[str, int, int, str]] = {}
    with path.open() as handle:
        for line in handle:
            if line.startswith("#") or not line.strip():
                continue
            parts = line.rstrip().split("\t")
            if len(parts) != 9 or parts[2] not in {"gene", "transcript", "mRNA"}:
                continue
            attrs = parts[8]
            match = re.search(r"(?:^|;)(?:geneID|ID)=([^;]+)", attrs)
            if not match:
                continue
            gene = strip_tx(match.group(1))
            if not gene.startswith("GY2U"):
                continue
            chrom, start, end, strand = parts[0], int(parts[3]), int(parts[4]), parts[6]
            if gene in coords:
                old = coords[gene]
                coords[gene] = (chrom, min(start, old[1]), max(end, old[2]), strand)
            else:
                coords[gene] = (chrom, start, end, strand)
    return coords


def parse_fasta_lengths(path: Path) -> dict[str, int]:
    lengths = {}
    name = None
    length = 0
    with path.open() as handle:
        for line in handle:
            if line.startswith(">"):
                if name is not None:
                    lengths[strip_tx(name)] = length
                name = line[1:].split()[0]
                length = 0
            else:
                length += len(line.strip())
        if name is not None:
            lengths[strip_tx(name)] = length
    return lengths


def plot_transition(summary: pd.DataFrame) -> None:
    fig, ax = plt.subplots(figsize=(7.2, 4.4))
    x = np.arange(len(summary))
    width = 0.36
    ax.bar(x - width / 2, summary["n_up"], width, color=ORANGE, label="Up-regulated")
    ax.bar(x + width / 2, summary["n_down"], width, color=BLUE, label="Down-regulated")
    ymax = summary[["n_up", "n_down"]].to_numpy().max()
    for i, row in summary.iterrows():
        ax.text(i, max(row["n_up"], row["n_down"]) + ymax * 0.035,
                f"Total {int(row['n_total_DEG']):,}", ha="center", va="bottom", fontsize=8)
    ax.set_xticks(x, summary["transition"])
    ax.set_ylabel("Number of DEGs")
    ax.set_xlabel("Adjacent developmental transition")
    ax.legend(frameon=False, ncol=2, loc="upper left")
    ax.spines[["top", "right"]].set_visible(False)
    ax.set_ylim(0, ymax * 1.22)
    ax.grid(axis="y", color=LIGHT, linewidth=0.6)
    fig.tight_layout()
    save_figure(fig, "Figure3D_adjacent_stage_transition")


def plot_enrichment(nonredundant: pd.DataFrame) -> None:
    sig = nonredundant[nonredundant["q_value"] < FDR].copy()
    if sig.empty:
        raise RuntimeError("No significant nonredundant GO/KEGG terms available for enrichment figure")
    sig["term_key"] = sig["database"] + " | " + sig["term_name"]
    best = sig.groupby("term_key")["q_value"].min().sort_values().head(18)
    keys = list(best.index)
    labels = []
    for key in keys:
        db, name = key.split(" | ", 1)
        labels.append(f"{db}: {name[:58]}")
    columns = [f"{STAGE_DAYS[a]}-{STAGE_DAYS[b]}" for _, a, b in TRANSITIONS]
    matrix = np.zeros((len(keys), len(columns)), dtype=float)
    for i, key in enumerate(keys):
        sub = sig[sig["term_key"] == key]
        for j, (contrast, _, _) in enumerate(TRANSITIONS):
            cand = sub[sub["transition"] == contrast]
            if cand.empty:
                continue
            row = cand.sort_values("q_value").iloc[0]
            val = min(-math.log10(max(float(row["q_value"]), 1e-300)), 20)
            matrix[i, j] = val if row["direction"] == "UP" else -val
    vmax = max(2.0, float(np.abs(matrix).max()))
    cmap = LinearSegmentedColormap.from_list("blue_white_orange", [BLUE, "#FFFFFF", ORANGE])
    fig, ax = plt.subplots(figsize=(8.3, max(5.0, 0.34 * len(keys) + 1.8)))
    image = ax.imshow(matrix, aspect="auto", cmap=cmap, vmin=-vmax, vmax=vmax)
    ax.set_xticks(np.arange(len(columns)), columns, rotation=22, ha="right")
    ax.set_yticks(np.arange(len(keys)), labels)
    ax.set_xlabel("Adjacent developmental transition (days after planting)")
    for x in np.arange(len(columns) + 1) - 0.5:
        ax.axvline(x, color="white", lw=0.7)
    for y in np.arange(len(keys) + 1) - 0.5:
        ax.axhline(y, color="white", lw=0.7)
    cb = fig.colorbar(image, ax=ax, fraction=0.03, pad=0.02)
    cb.set_label("Signed -log10(q): UP (+), DOWN (-)")
    fig.tight_layout()
    save_figure(fig, "FigureS_transition_functional_enrichment")


def module_color(name: str) -> str:
    aliases = {"brown": "#A65628", "yellow": "#E6AB02", "green": "#1B9E77",
               "red": "#D55E00", "blue": "#377EB8", "turquoise": "#40B0A6",
               "black": "#222222", "pink": "#E78AC3", "magenta": "#CC79A7",
               "purple": "#984EA3", "greenyellow": "#A6D854", "tan": "#C49A6C",
               "salmon": "#FC8D62", "cyan": "#56B4E9", "grey": "#9CA3AF"}
    return aliases.get(str(name), "#777777")


def plot_wgcna(integration: pd.DataFrame) -> None:
    data = integration.sort_values("order").copy()
    labels = [f"{s} ({g})" for s, g in zip(data["symbol"], data["gene_id"])]
    y = np.arange(len(data))
    fig, (ax, ax2) = plt.subplots(1, 2, figsize=(11.2, 7.4),
                                  gridspec_kw={"width_ratios": [2.4, 1.15], "wspace": 0.12})
    colors = [module_color(x) for x in data["WGCNA_module"]]
    ax.axvline(0, color=LIGHT, lw=1)
    ax.axvline(0.8, color=MUTED, lw=0.8, ls="--")
    ax.axvline(-0.8, color=MUTED, lw=0.8, ls="--")
    for yi, value, color in zip(y, data["kME"], colors):
        if pd.notna(value):
            ax.plot([0, value], [yi, yi], color=color, lw=1.5, alpha=0.75)
            ax.scatter(value, yi, s=42, color=color, edgecolor="white", linewidth=0.5, zorder=3)
    ax.set_yticks(y, labels, fontstyle="italic", fontsize=7.5)
    ax.set_xlim(-1.05, 1.05)
    ax.set_xlabel("Module membership (kME)")
    ax.set_ylim(len(data) - 0.5, -0.5)
    ax.spines[["top", "right", "left"]].set_visible(False)
    ax.tick_params(axis="y", length=0)
    stage_numeric = data["module_peak_stage"].str.extract(r"(\d)")[0].astype(float)
    flags = np.column_stack([
        stage_numeric,
        data["strict_corm_enriched"].astype(int),
        (data["development_LRT_q"] < FDR).astype(int),
    ])
    cmaps = [plt.cm.viridis, ListedColormap(["#F3F4F6", GREEN]), ListedColormap(["#F3F4F6", ORANGE])]
    for col in range(3):
        vals = flags[:, col][:, None]
        if col == 0:
            ax2.imshow(vals, extent=(col - 0.42, col + 0.42, len(data) - 0.5, -0.5),
                       aspect="auto", cmap=cmaps[col], vmin=1, vmax=6)
        else:
            ax2.imshow(vals, extent=(col - 0.42, col + 0.42, len(data) - 0.5, -0.5),
                       aspect="auto", cmap=cmaps[col], vmin=0, vmax=1)
        for yi, value in enumerate(flags[:, col]):
            text = f"S{int(value)}" if col == 0 and np.isfinite(value) else ("Yes" if value == 1 else "No")
            ax2.text(col, yi, text, ha="center", va="center", fontsize=6.8,
                     color="white" if (col == 0 and value >= 4) or (col > 0 and value == 1) else INK)
    ax2.set_xlim(-0.5, 2.5)
    ax2.set_ylim(len(data) - 0.5, -0.5)
    ax2.set_xticks([0, 1, 2], ["Module\npeak", "Strict corm\nenriched", "Development\nresponsive"])
    ax2.set_yticks([])
    ax2.spines[:].set_visible(False)
    fig.tight_layout()
    save_figure(fig, "FigureS_starch_gene_module_integration")


def plot_conservation(conservation: pd.DataFrame) -> None:
    status_order = ["reference", "1:1 syntenic ortholog", "syntenic duplicated ortholog",
                    "non-syntenic homolog", "annotation-missing but genomic sequence recovered",
                    "candidate true absence", "unresolved"]
    colors = ["#4D4D4D", GREEN, SKY, YELLOW, PURPLE, ORANGE, "#D1D5DB"]
    code = {name: i for i, name in enumerate(status_order)}
    rows = conservation.sort_values("order").copy()
    matrix = []
    annotations = []
    for _, row in rows.iterrows():
        matrix.append([code["reference"], code.get(row["LXY_status"], code["unresolved"]),
                       code.get(row["BunLong_status"], code["unresolved"])])
        annotations.append([f"n={int(row['GY2_copy_number'])}" if pd.notna(row["GY2_copy_number"]) else "n=?",
                            f"n={int(row['LXY_copy_number'])}" if pd.notna(row["LXY_copy_number"]) else "n=?",
                            f"n={int(row['BunLong_copy_number'])}" if pd.notna(row["BunLong_copy_number"]) else "n=?"])
    labels = [f"{s} ({g})" for s, g in zip(rows["symbol"], rows["GY2_gene_id"])]
    fig, ax = plt.subplots(figsize=(8.7, 7.2))
    ax.imshow(np.array(matrix), aspect="auto", cmap=ListedColormap(colors), vmin=-0.5,
              vmax=len(colors) - 0.5)
    ax.set_xticks([0, 1, 2], ["GY2", "LXY", "Bun Long"])
    ax.set_yticks(np.arange(len(rows)), labels, fontstyle="italic", fontsize=7.5)
    for i in range(len(rows)):
        for j in range(3):
            ax.text(j, i, annotations[i][j], ha="center", va="center", fontsize=7,
                    color="white" if matrix[i][j] in {0, 1, 4, 5} else INK)
    ax.set_xticks(np.arange(-0.5, 3, 1), minor=True)
    ax.set_yticks(np.arange(-0.5, len(rows), 1), minor=True)
    ax.grid(which="minor", color="white", linewidth=1)
    ax.tick_params(which="minor", bottom=False, left=False)
    handles = [plt.Rectangle((0, 0), 1, 1, color=colors[code[s]]) for s in status_order[:4] + ["unresolved"]]
    legend_labels = status_order[:4] + ["unresolved"]
    ax.legend(handles, legend_labels, frameon=False, bbox_to_anchor=(1.02, 1), loc="upper left", fontsize=7)
    fig.tight_layout()
    save_figure(fig, "FigureS_three_genome_starch_gene_conservation")


def main() -> None:
    ensure_dirs()

    # Input paths.
    retained_dev_dir = INPUT / "expression" / "deseq2" / "PRJNA756190"
    dev_meta_path = INPUT / "expression" / "developmental_sample_map.tsv"
    dev_counts_path = INPUT / "expression" / "counts" / "PRJNA756190.gene_counts.tsv"
    dev_dir = INPUT / "developmental" / "results"
    tis_dir = INPUT / "expression" / "deseq2" / "PRJNA1493089"
    tis_counts_path = INPUT / "expression" / "counts" / "PRJNA1493089.gene_counts.tsv"
    recompute_r_path = INPUT / "developmental" / ".." / ".." / "scripts" / "developmental_expression.R"
    recompute_expected_sha_path = INPUT / "developmental" / "input_checksums.sha256"
    contrast_dir = dev_dir / "contrasts"
    selected_path = INPUT / "pathway" / "Figure4_selected_genes.tsv"
    catalog_path = INPUT / "pathway" / "starch_sucrose_candidate_gene_catalog.tsv"
    id_map_path = INPUT / "annotation" / "GY2.v3.author_id_mapping.tsv"
    gff_path = INPUT / "annotation" / "GY2.uniform_reannotated.v3.gff3"
    pep_path = INPUT / "annotation" / "GY2.uniform_reannotated.v3.primary.pep.fa"
    og_path = INPUT / "comparative" / "Orthogroups.tsv"
    lxy_anchor_path = INPUT / "comparative" / "GY2_vs_LXY.anchor_pairs.tsv"
    bun_anchor_path = INPUT / "comparative" / "GY2_vs_Bun_Long.anchor_pairs.tsv"
    go_path = INPUT / "functional_annotation" / "Cesc.pep.filter.GO.anno.txt"
    kegg_path = INPUT / "functional_annotation" / "Cesc.pep.filter.Kegg.pathway"
    supported_path = INPUT / "functional_annotation" / "GY2_starch_sucrose_supported_genes.tsv"
    wgcna_assign_path = dev_dir / "WGCNA_module_assignments.tsv"
    wgcna_me_path = dev_dir / "WGCNA_module_eigengenes.tsv"
    wgcna_stage_path = dev_dir / "WGCNA_module_stage_summary.tsv"

    required = [dev_dir / "vst_expression.tsv.gz", dev_meta_path, dev_counts_path,
                dev_dir / "omnibus_LRT_all_genes.tsv", tis_dir / "vst_expression.tsv.gz",
                tis_dir / "sample_metadata.tsv", tis_counts_path,
                tis_dir / "omnibus_LRT_all_genes.tsv",
                selected_path, catalog_path, id_map_path, gff_path, pep_path, og_path,
                lxy_anchor_path, bun_anchor_path, go_path, kegg_path, supported_path,
                wgcna_assign_path, wgcna_me_path, wgcna_stage_path,
                dev_dir / "developmental_summary.json",
                dev_dir / "feature_filter.tsv", dev_dir / "WGCNA_soft_threshold.tsv",
                dev_dir / "sessionInfo.txt",
                INPUT / "developmental" / "analysis_metadata.tsv",
                recompute_r_path, recompute_expected_sha_path,
                retained_dev_dir / "vst_expression.tsv.gz",
                retained_dev_dir / "omnibus_LRT_all_genes.tsv"]
    for contrast, _, _ in TRANSITIONS:
        required.append(contrast_dir / f"{contrast}.all_genes.tsv")
    for tissue in ("leaf", "petiole", "root"):
        required.append(tis_dir / "contrasts" / f"{tissue}_vs_corm.all_genes.tsv")
    missing = [str(p) for p in required if not p.exists()]
    if missing:
        raise FileNotFoundError("Missing required inputs:\n" + "\n".join(missing))

    selected = read_tsv(selected_path)
    catalog = read_tsv(catalog_path)
    if len(selected) != 20 or selected["gene_id"].nunique() != 20:
        raise RuntimeError("Figure4_selected_genes.tsv must contain exactly 20 unique final-v3 genes")
    selected_ids = set(selected["gene_id"])

    dev_meta = read_tsv(dev_meta_path)
    tis_meta = read_tsv(tis_dir / "sample_metadata.tsv")
    dev_vst = read_tsv(dev_dir / "vst_expression.tsv.gz").set_index("gene_id")
    tis_vst = read_tsv(tis_dir / "vst_expression.tsv.gz").set_index("gene_id")
    dev_samples = list(dev_vst.columns)
    tis_samples = list(tis_vst.columns)
    if set(dev_meta["sample_id"]) != set(dev_samples) or set(tis_meta.iloc[:, 0]) != set(tis_samples):
        raise RuntimeError("Sample IDs do not match VST matrices")
    if dev_meta.groupby("stage").size().to_dict() != {s: 3 for s in STAGES}:
        raise RuntimeError("Developmental biological replicate counts are inconsistent")
    if tis_meta.groupby("tissue").size().to_dict() != {x: 3 for x in ["corm", "leaf", "petiole", "root"]}:
        raise RuntimeError("Tissue biological replicate counts are inconsistent")
    if not selected_ids <= set(dev_vst.index) or not selected_ids <= set(tis_vst.index):
        raise RuntimeError("One or more Figure 4 genes are missing from final-v3 expression matrices")

    # Compare developmental expression matrices.
    retained_dev_vst = read_tsv(retained_dev_dir / "vst_expression.tsv.gz").set_index("gene_id")
    if set(retained_dev_vst.index) != set(dev_vst.index) or set(retained_dev_vst.columns) != set(dev_vst.columns):
        raise RuntimeError("Fresh and retained final-v3-full VST universes differ")
    retained_dev_vst = retained_dev_vst.reindex(index=dev_vst.index, columns=dev_vst.columns)
    vst_abs_delta = np.abs(dev_vst.to_numpy(dtype=float) - retained_dev_vst.to_numpy(dtype=float))
    vst_max_abs_delta = float(np.nanmax(vst_abs_delta))
    vst_mean_abs_delta = float(np.nanmean(vst_abs_delta))
    vst_flat_correlation = float(np.corrcoef(
        dev_vst.to_numpy(dtype=float).ravel(), retained_dev_vst.to_numpy(dtype=float).ravel())[0, 1])
    if vst_max_abs_delta > 1e-10 or vst_flat_correlation < 0.999999999999:
        raise RuntimeError(
            "Fresh final-v3-full VST failed concordance thresholds: "
            f"max_abs={vst_max_abs_delta}, r={vst_flat_correlation}")

    # Gene identifier mapping.
    id_map = read_tsv(id_map_path)
    if id_map["v3_gene_id"].duplicated().any():
        raise RuntimeError("Final-v3 gene IDs are not unique in author ID mapping")
    mapped = id_map[id_map["author_gene_id"].notna() & (id_map["author_gene_id"] != ".")].copy()
    if mapped["author_gene_id"].duplicated().any():
        raise RuntimeError("Non-null author gene IDs are not one-to-one")
    author_to_v3 = dict(zip(mapped["author_gene_id"], mapped["v3_gene_id"]))
    v3_to_author = dict(zip(mapped["v3_gene_id"], mapped["author_gene_id"]))

    # Analysis 1: adjacent-stage DEG transition strength.
    transition_rows = []
    foregrounds: dict[tuple[str, str], set[str]] = {}
    backgrounds: dict[str, set[str]] = {}
    for contrast, stage_from, stage_to in TRANSITIONS:
        data = read_tsv(contrast_dir / f"{contrast}.all_genes.tsv")
        if data["gene_id"].duplicated().any():
            raise RuntimeError(f"Duplicate gene IDs in {contrast}")
        qcol = "padj_BH_all_eligible"
        tested = data[data["pvalue"].notna() & np.isfinite(data["pvalue"])].copy()
        sig = tested[(tested[qcol] < FDR) & (tested["log2FoldChange"].abs() >= LFC)]
        up = set(sig.loc[sig["log2FoldChange"] >= LFC, "gene_id"])
        down = set(sig.loc[sig["log2FoldChange"] <= -LFC, "gene_id"])
        backgrounds[contrast] = set(tested["gene_id"])
        foregrounds[(contrast, "UP")] = up
        foregrounds[(contrast, "DOWN")] = down
        transition_rows.append({
            "transition": f"{STAGE_DAYS[stage_from]}-{STAGE_DAYS[stage_to]} d",
            "contrast_id": contrast,
            "stage_from": stage_from,
            "stage_to": stage_to,
            "n_tested": len(tested),
            "n_up": len(up),
            "n_down": len(down),
            "n_total_DEG": len(up) + len(down),
            "threshold_FDR": FDR,
            "threshold_log2FC": LFC,
        })
    transition_summary = pd.DataFrame(transition_rows)
    write_tsv(transition_summary, TABLES / "01_Adjacent_stage_DEG_transition_summary.tsv")

    # Analysis 2: GO BP and KEGG ORA with contrast-specific tested backgrounds.
    go_terms, go_meta = parse_go(go_path, author_to_v3)
    go_terms = {k: v for k, v in go_terms.items() if go_meta[k]["ontology"] == "Biological Process"}
    kegg_terms, kegg_meta = parse_kegg(kegg_path, author_to_v3)
    go_results = []
    kegg_results = []
    for contrast, _, _ in TRANSITIONS:
        for direction in ("UP", "DOWN"):
            go_results.append(enrich_one(foregrounds[(contrast, direction)], backgrounds[contrast],
                                         go_terms, go_meta, contrast, direction, "GO_BP"))
            kegg_results.append(enrich_one(foregrounds[(contrast, direction)], backgrounds[contrast],
                                           kegg_terms, kegg_meta, contrast, direction, "KEGG"))
    go_enrichment = pd.concat(go_results, ignore_index=True) if go_results else pd.DataFrame()
    kegg_enrichment = pd.concat(kegg_results, ignore_index=True) if kegg_results else pd.DataFrame()
    write_tsv(go_enrichment, TABLES / "02_Adjacent_stage_GO_enrichment.tsv")
    write_tsv(kegg_enrichment, TABLES / "03_Adjacent_stage_KEGG_enrichment.tsv")
    nonredundant = reduce_redundancy(go_enrichment, kegg_enrichment)
    write_tsv(nonredundant, TABLES / "Adjacent_stage_enrichment_nonredundant.tsv")

    # Analysis 3: invert retained X-vs-corm effects to obtain corm-vs-X effects.
    tissue_all = None
    for tissue in ("leaf", "petiole", "root"):
        data = read_tsv(tis_dir / "contrasts" / f"{tissue}_vs_corm.all_genes.tsv")
        part = data[["gene_id", "log2FoldChange", "padj"]].copy()
        part[f"corm_vs_{tissue}_log2FC"] = -part.pop("log2FoldChange")
        part[f"corm_vs_{tissue}_q"] = part.pop("padj")
        tissue_all = part if tissue_all is None else tissue_all.merge(part, on="gene_id", how="inner", validate="one_to_one")
    symbol_map = dict(zip(catalog["gene_id"], catalog["symbol"]))
    tissue_all.insert(1, "symbol", tissue_all["gene_id"].map(symbol_map))
    strict = np.ones(len(tissue_all), dtype=bool)
    positive = np.ones(len(tissue_all), dtype=bool)
    sig_count = np.zeros(len(tissue_all), dtype=int)
    for tissue in ("leaf", "petiole", "root"):
        lfc = tissue_all[f"corm_vs_{tissue}_log2FC"]
        q = tissue_all[f"corm_vs_{tissue}_q"]
        strict &= (lfc > LFC) & (q < FDR)
        positive &= lfc > 0
        sig_count += (q < FDR).fillna(False).astype(int).to_numpy()
    relaxed = positive & (sig_count >= 2) & ~strict
    tissue_all["strict_corm_enriched"] = strict
    tissue_all["relaxed_corm_biased"] = relaxed
    tissue_all["classification"] = np.where(strict, "strict corm-enriched",
                                               np.where(relaxed, "corm-biased relaxed", "not classified"))
    write_tsv(tissue_all, TABLES / "Tissue_DEG_all_contrasts.tsv")
    write_tsv(tissue_all[tissue_all["strict_corm_enriched"]].copy(), TABLES / "04_Corm_enriched_strict.tsv")
    write_tsv(tissue_all[tissue_all["relaxed_corm_biased"]].copy(), TABLES / "05_Corm_biased_relaxed.tsv")

    # Analysis 4: developmental-responsive x strict corm-enriched intersection.
    dev_lrt = read_tsv(dev_dir / "omnibus_LRT_all_genes.tsv")
    if "padj_BH_all_eligible" in dev_lrt.columns:
        dev_lrt = dev_lrt.rename(columns={"padj_BH_all_eligible": "padj"})
    if "padj" not in dev_lrt.columns:
        raise RuntimeError("Developmental LRT lacks a BH-adjusted q-value column")
    retained_dev_lrt = read_tsv(retained_dev_dir / "omnibus_LRT_all_genes.tsv")
    if "padj_BH_all_eligible" in retained_dev_lrt.columns:
        retained_dev_lrt = retained_dev_lrt.rename(columns={"padj_BH_all_eligible": "padj"})
    if set(retained_dev_lrt["gene_id"]) != set(dev_lrt["gene_id"]):
        raise RuntimeError("Fresh and retained final-v3-full LRT gene universes differ")
    fresh_lrt_audit = dev_lrt.set_index("gene_id").sort_index()
    retained_lrt_audit = retained_dev_lrt.set_index("gene_id").sort_index()
    lrt_numeric_columns = ["baseMean", "log2FoldChange", "lfcSE", "stat", "pvalue", "padj"]
    lrt_max_abs_deltas = {
        column: float(np.nanmax(np.abs(
            fresh_lrt_audit[column].to_numpy(dtype=float) -
            retained_lrt_audit[column].to_numpy(dtype=float))))
        for column in lrt_numeric_columns
    }
    lrt_finite_masks_equal = {
        column: bool(np.array_equal(
            np.isfinite(fresh_lrt_audit[column].to_numpy(dtype=float)),
            np.isfinite(retained_lrt_audit[column].to_numpy(dtype=float))))
        for column in lrt_numeric_columns
    }
    lrt_bh05_agreement = float(np.mean(
        (fresh_lrt_audit["padj"] < FDR).to_numpy() ==
        (retained_lrt_audit["padj"] < FDR).to_numpy()))
    fresh_dynamic_bh05 = int((fresh_lrt_audit["padj"] < FDR).sum())
    retained_dynamic_bh05 = int((retained_lrt_audit["padj"] < FDR).sum())
    if (not all(lrt_finite_masks_equal.values()) or lrt_bh05_agreement != 1.0 or
            fresh_dynamic_bh05 != 11884 or retained_dynamic_bh05 != 11884):
        raise RuntimeError(
            "Fresh final-v3-full LRT failed concordance thresholds: "
            f"finite_masks={lrt_finite_masks_equal}, agreement={lrt_bh05_agreement}, "
            f"dynamic={fresh_dynamic_bh05}/{retained_dynamic_bh05}")
    concordance_audit = {
        "status": "PASS",
        "comparison": "fresh final-v3-full recomputation versus retained task109 final-v3-full outputs",
        "vst_gene_ids_equal": True,
        "vst_sample_ids_equal": True,
        "vst_max_absolute_difference": vst_max_abs_delta,
        "vst_mean_absolute_difference": vst_mean_abs_delta,
        "vst_flattened_Pearson_r": vst_flat_correlation,
        "lrt_gene_ids_equal": True,
        "lrt_finite_value_masks_equal": lrt_finite_masks_equal,
        "lrt_max_absolute_differences": lrt_max_abs_deltas,
        "lrt_BH05_classification_agreement": lrt_bh05_agreement,
        "fresh_dynamic_BH05": fresh_dynamic_bh05,
        "retained_dynamic_BH05": retained_dynamic_bh05,
    }
    (QC / "FINAL_V3_FULL_RECOMPUTE_CONCORDANCE.json").write_text(
        json.dumps(concordance_audit, indent=2) + "\n", encoding="utf-8")
    dev_lrt = dev_lrt[["gene_id", "baseMean", "stat", "pvalue", "padj"]].rename(
        columns={"baseMean": "development_baseMean", "stat": "development_LRT_stat",
                 "pvalue": "development_LRT_p", "padj": "development_LRT_q"})
    dev_lrt["development_responsive"] = dev_lrt["development_LRT_q"] < FDR
    merged_dev_tissue = tissue_all.merge(dev_lrt, on="gene_id", how="inner", validate="one_to_one")
    merged_dev_tissue["corm_developmental"] = (merged_dev_tissue["strict_corm_enriched"] &
                                                 merged_dev_tissue["development_responsive"])
    corm_dev = merged_dev_tissue[merged_dev_tissue["corm_developmental"]].copy()
    write_tsv(corm_dev, TABLES / "06_Corm_developmental_gene_set.tsv")

    universe = set(merged_dev_tissue["gene_id"])
    cd_set = set(corm_dev["gene_id"])
    pathway_set = selected_ids & universe
    a = len(cd_set & pathway_set)
    b = len(pathway_set - cd_set)
    c = len(cd_set - pathway_set)
    d = len(universe - cd_set - pathway_set)
    pathway_or, pathway_p = fisher_exact([[a, b], [c, d]], alternative="greater")
    corrected_or, ci_low, ci_high = odds_ci(a, b, c, d)
    pathway_test = pd.DataFrame([{
        "gene_set": "Figure4_starch_sucrose_20",
        "universe_n": len(universe),
        "gene_set_n": len(pathway_set),
        "corm_developmental_n": len(cd_set),
        "overlap_n": a,
        "odds_ratio_fisher": pathway_or,
        "odds_ratio_Haldane_corrected": corrected_or,
        "odds_ratio_95CI_low": ci_low,
        "odds_ratio_95CI_high": ci_high,
        "p_value": pathway_p,
        "q_value": pathway_p,
    }])
    write_tsv(pathway_test, TABLES / "Corm_developmental_starch_enrichment.tsv")

    # Analysis 5: recompute kME using exact 18-sample alignment, without rebuilding WGCNA.
    assignments = read_tsv(wgcna_assign_path)
    eigengenes = read_tsv(wgcna_me_path).set_index("sample_id")
    stage_summary = read_tsv(wgcna_stage_path)
    if assignments["gene_id"].duplicated().any():
        raise RuntimeError("WGCNA assignment gene IDs are not unique")
    dev_order = []
    for sid in eigengenes.index:
        current = re.fullmatch(r"DEV_S(\d+)_R(\d+)", sid)
        legacy = re.fullmatch(r"S(\d+)-(\d+)", sid)
        if current:
            dev_order.append(sid)
        elif legacy:
            dev_order.append(f"DEV_S{legacy.group(1)}_R{legacy.group(2)}")
        else:
            raise RuntimeError(f"Unrecognized WGCNA sample ID: {sid}")
    if set(dev_order) != set(dev_vst.columns):
        raise RuntimeError("WGCNA eigengene sample order cannot be reconciled with final-v3 VST")
    expr = dev_vst[dev_order]
    assignment_genes = set(assignments["gene_id"])
    vst_genes = set(expr.index)
    assignment_not_in_vst = assignment_genes - vst_genes
    vst_without_assignment = vst_genes - assignment_genes
    assigned_in_vst = assignment_genes & vst_genes

    def count_zero_mad(genes: set[str]) -> int:
        if not genes:
            return 0
        values = expr.loc[sorted(genes)].to_numpy(dtype=float)
        medians = np.median(values, axis=1, keepdims=True)
        mad = np.median(np.abs(values - medians), axis=1)
        return int(np.isclose(mad, 0.0, atol=1e-12).sum())

    vst_without_assignment_zero_mad = count_zero_mad(vst_without_assignment)
    assigned_in_vst_zero_mad = count_zero_mad(assigned_in_vst)
    kme_rows = []
    for _, row in assignments.iterrows():
        gene = row["gene_id"]
        module = row["module_id"]
        if module == "grey" or gene not in expr.index or f"ME{module}" not in eigengenes.columns:
            continue
        x = expr.loc[gene].to_numpy(dtype=float)
        y = eigengenes[f"ME{module}"].to_numpy(dtype=float)
        if np.std(x) == 0 or np.std(y) == 0:
            r, p = np.nan, np.nan
        else:
            r, p = pearsonr(x, y)
        kme_rows.append({"gene_id": gene, "module_id": module, "kME": r, "kME_p": p,
                         "retained_own_module_kME": row.get("own_module_kME", np.nan)})
    all_kme = pd.DataFrame(kme_rows)
    all_kme["kME_q"] = bh_adjust(all_kme["kME_p"])
    all_kme["absolute_kME"] = all_kme["kME"].abs()
    all_kme["high_module_membership"] = (all_kme["absolute_kME"] >= 0.80) & (all_kme["kME_q"] < FDR)
    write_tsv(all_kme, TABLES / "WGCNA_all_gene_kME.tsv")

    module_sizes = assignments.groupby("module_id").size().rename("module_size")
    stage_info = stage_summary.set_index("module_id")
    wgcna = selected[["order", "gene_id", "symbol", "category"]].copy()
    wgcna = wgcna.merge(assignments[["gene_id", "module_id"]], on="gene_id", how="left", validate="one_to_one")
    wgcna = wgcna.merge(all_kme.drop(columns="module_id"), on="gene_id", how="left", validate="one_to_one")
    wgcna["module_size"] = wgcna["module_id"].map(module_sizes)
    wgcna["module_stage_rho"] = wgcna["module_id"].map(stage_info["stage_mean_Spearman_rho"])
    wgcna["module_peak_stage"] = wgcna["module_id"].map(stage_info["peak_stage"])
    wgcna = wgcna.rename(columns={"category": "pathway_category", "module_id": "WGCNA_module"})
    wgcna = wgcna.merge(merged_dev_tissue[["gene_id", "development_LRT_q", "strict_corm_enriched",
                                           "relaxed_corm_biased"]], on="gene_id", how="left", validate="one_to_one")
    write_tsv(wgcna, TABLES / "07_Starch_genes_WGCNA_integration.tsv")

    eligible = set(assignments["gene_id"])
    pathway_eligible = selected_ids & eligible
    module_rows = []
    for module, group in assignments[assignments["module_id"] != "grey"].groupby("module_id"):
        module_set = set(group["gene_id"])
        aa = len(pathway_eligible & module_set)
        bb = len(pathway_eligible - module_set)
        cc = len(module_set - pathway_eligible)
        dd = len(eligible - module_set - pathway_eligible)
        odds, p = fisher_exact([[aa, bb], [cc, dd]], alternative="greater")
        module_rows.append({"module": module, "module_n": len(module_set),
                            "pathway_genes_in_module": aa,
                            "pathway_gene_ids": ";".join(sorted(pathway_eligible & module_set)),
                            "odds_ratio": odds, "p_value": p})
    module_enrich = pd.DataFrame(module_rows)
    module_enrich["q_value"] = bh_adjust(module_enrich["p_value"])
    module_enrich = module_enrich.sort_values(["q_value", "p_value", "module"])
    write_tsv(module_enrich, TABLES / "08_WGCNA_starch_module_enrichment.tsv")

    # Analysis 6: final-v3 orthogroup + MCScanX synteny status.
    coords = parse_gff_coordinates(gff_path)
    protein_lengths = parse_fasta_lengths(pep_path)
    orthogroups = read_tsv(og_path)
    gy2_to_og = {}
    og_members = {}
    for _, row in orthogroups.iterrows():
        og = row["Orthogroup"]
        per_species = {sp: split_members(row.get(sp, "")) for sp in ["GY2", "LXY", "Bun_Long"]}
        og_members[og] = per_species
        for member in per_species["GY2"]:
            gene = strip_tx(member)
            if gene in gy2_to_og:
                raise RuntimeError(f"GY2 gene maps to multiple orthogroups: {gene}")
            gy2_to_og[gene] = og

    def anchor_map(path: Path) -> tuple[dict[str, list[str]], dict[str, list[str]]]:
        data = read_tsv(path)
        targets = defaultdict(list)
        scores = defaultdict(list)
        for _, row in data.iterrows():
            left = strip_tx(row["left_gene"])
            right = strip_tx(row["right_gene"])
            targets[left].append(right)
            scores[left].append(str(row["blast_evalue_or_score"]))
        return {k: sorted(set(v)) for k, v in targets.items()}, scores

    lxy_anchor, lxy_anchor_score = anchor_map(lxy_anchor_path)
    bun_anchor, bun_anchor_score = anchor_map(bun_anchor_path)

    def comparator_status(gene: str, species: str, anchors: dict[str, list[str]], og: str | None):
        members = og_members.get(og, {}).get(species, []) if og else []
        members_clean = sorted(set(strip_tx(x) for x in members))
        anchored = sorted(set(anchors.get(gene, [])))
        anchored_in_og = sorted(set(anchored) & set(members_clean))
        extra_anchors = sorted(set(anchored) - set(members_clean))
        gy2_count = len(og_members.get(og, {}).get("GY2", [])) if og else 0
        if anchored_in_og:
            strict_one_to_one = (
                gy2_count == 1
                and len(members_clean) == 1
                and len(anchored_in_og) == 1
                and not extra_anchors
            )
            status = "1:1 syntenic ortholog" if strict_one_to_one else "syntenic duplicated ortholog"
            genes = anchored
        elif members_clean:
            status = "non-syntenic homolog"
            genes = members_clean
        else:
            status = "unresolved"
            genes = []
        return status, genes, len(members_clean), bool(anchored_in_og), extra_anchors

    conservation_rows = []
    for _, row in selected.sort_values("order").iterrows():
        gene = row["gene_id"]
        og = gy2_to_og.get(gene)
        if og is None:
            raise RuntimeError(f"Final-v3 Figure 4 gene has no unique Orthogroup mapping: {gene}")
        l_status, l_genes, l_count, l_syn, l_extra = comparator_status(gene, "LXY", lxy_anchor, og)
        b_status, b_genes, b_count, b_syn, b_extra = comparator_status(gene, "Bun_Long", bun_anchor, og)
        if l_status == "1:1 syntenic ortholog" and b_status == "1:1 syntenic ortholog":
            combined_status = "1:1 syntenic ortholog"
        elif "syntenic duplicated ortholog" in {l_status, b_status}:
            combined_status = "syntenic duplicated ortholog"
        elif "non-syntenic homolog" in {l_status, b_status}:
            combined_status = "non-syntenic homolog"
        else:
            combined_status = "unresolved"
        chrom, start, end, strand = coords.get(gene, (np.nan, np.nan, np.nan, np.nan))
        conservation_rows.append({
            "order": row["order"], "GY2_gene_id": gene, "symbol": row["symbol"],
            "pathway": row["category"], "orthogroup": og, "GY2_chr": chrom,
            "GY2_start": start, "GY2_end": end, "GY2_strand": strand,
            "GY2_copy_number": len(og_members[og]["GY2"]),
            "LXY_gene": ";".join(l_genes), "LXY_status": l_status, "LXY_syntenic": l_syn,
            "LXY_copy_number": l_count, "LXY_protein_identity": np.nan,
            "LXY_protein_coverage": np.nan,
            "LXY_extra_cross_orthogroup_anchors": ";".join(l_extra),
            "BunLong_gene": ";".join(b_genes), "BunLong_status": b_status,
            "BunLong_syntenic": b_syn, "BunLong_copy_number": b_count,
            "BunLong_protein_identity": np.nan, "BunLong_protein_coverage": np.nan,
            "BunLong_extra_cross_orthogroup_anchors": ";".join(b_extra),
            "combined_status": combined_status,
        })
    conservation = pd.DataFrame(conservation_rows)
    write_tsv(conservation, TABLES / "09_Starch_pathway_three_genome_conservation.tsv")

    # Functional annotation QC for the fixed 20-gene set.
    supported = read_tsv(supported_path)
    supported = supported.rename(columns={"gene_id": "original_gene_id"})
    qc_annot = selected.merge(catalog[["gene_id", "original_gene_id", "description", "annotation_source"]],
                              on="gene_id", how="left", suffixes=("", "_catalog"), validate="one_to_one")
    keep_cols = [c for c in ["original_gene_id", "GO_annotation", "KEGG_annotation", "Pfam_annotation",
                              "Swissprot_annotation", "nr_annotation", "evidence_status"] if c in supported.columns]
    qc_annot = qc_annot.merge(supported[keep_cols], on="original_gene_id", how="left", validate="many_to_one")
    qc_annot["protein_length_aa"] = qc_annot["gene_id"].map(protein_lengths)
    qc_annot["best_protein_hit"] = qc_annot.get("Swissprot_annotation", qc_annot["description"]).fillna(qc_annot["description"])
    qc_annot["hit_species"] = qc_annot["best_protein_hit"].str.extract(r"OS=(.*?)\s+OX=")[0]
    qc_annot["hit_identity"] = np.nan
    qc_annot["hit_coverage"] = np.nan
    qc_annot["orthogroup"] = qc_annot["gene_id"].map(gy2_to_og)
    qc_annot["LXY_syntenic"] = qc_annot["gene_id"].isin(lxy_anchor)
    qc_annot["BunLong_syntenic"] = qc_annot["gene_id"].isin(bun_anchor)
    qc_annot["coordinate_present"] = qc_annot["gene_id"].isin(coords)
    robust = (qc_annot["protein_length_aa"].fillna(0) > 0) & qc_annot["coordinate_present"] & qc_annot["orthogroup"].notna()
    has_annotation = qc_annot["description"].notna() & qc_annot["annotation_source"].notna()
    qc_annot["annotation_confidence"] = np.where(robust & has_annotation & (qc_annot["LXY_syntenic"] | qc_annot["BunLong_syntenic"]),
                                                   "moderate-to-high homology-supported",
                                                   np.where(robust & has_annotation, "moderate homology-supported", "ambiguous"))
    functional_qc = qc_annot.rename(columns={"gene_id": "gene_id", "symbol": "current_symbol"})
    write_tsv(functional_qc, TABLES / "12_Starch_gene_annotation_evidence.tsv")

    # Required synthesis evidence matrix.
    stage_means = {}
    for gene in selected_ids:
        stage_means[gene] = {}
        for stage in STAGES:
            cols = [c for c in dev_vst.columns if c.startswith(f"DEV_{stage}_")]
            stage_means[gene][stage] = float(dev_vst.loc[gene, cols].mean())
    evidence = selected[["order", "gene_id", "symbol", "category"]].rename(columns={"category": "pathway_category"})
    evidence = evidence.merge(conservation.drop(columns=["order", "symbol", "pathway"]),
                              left_on="gene_id", right_on="GY2_gene_id", how="left", validate="one_to_one")
    evidence = evidence.drop(columns="GY2_gene_id")
    evidence = evidence.rename(columns={"GY2_chr": "chr", "GY2_start": "start", "GY2_end": "end",
                                        "GY2_copy_number": "copy_number_GY2", "LXY_copy_number": "copy_number_LXY",
                                        "BunLong_copy_number": "copy_number_BunLong"})
    evidence = evidence.merge(dev_lrt[["gene_id", "development_LRT_q"]], on="gene_id", how="left", validate="one_to_one")
    evidence["development_peak_stage"] = evidence["gene_id"].map(
        lambda g: max(stage_means[g], key=stage_means[g].get))
    evidence["development_expression_amplitude"] = evidence["gene_id"].map(
        lambda g: max(stage_means[g].values()) - min(stage_means[g].values()))
    evidence = evidence.merge(wgcna[["gene_id", "WGCNA_module", "module_stage_rho", "module_peak_stage", "kME", "kME_q"]],
                              on="gene_id", how="left", validate="one_to_one")
    tissue_cols = ["gene_id"] + [c for c in tissue_all.columns if c.startswith("corm_vs_") or c in
                                  {"strict_corm_enriched", "relaxed_corm_biased"}]
    evidence = evidence.merge(tissue_all[tissue_cols], on="gene_id", how="left", validate="one_to_one")
    evidence["corm_developmental"] = ((evidence["development_LRT_q"] < FDR) & evidence["strict_corm_enriched"])
    conf_map = dict(zip(functional_qc["gene_id"], functional_qc["annotation_confidence"]))
    evidence["annotation_confidence"] = evidence["gene_id"].map(conf_map)
    evidence["criterion_development_responsive"] = evidence["development_LRT_q"] < FDR
    evidence["criterion_strict_corm_enriched"] = evidence["strict_corm_enriched"]
    evidence["criterion_non_grey_module"] = evidence["WGCNA_module"].notna() & (evidence["WGCNA_module"] != "grey")
    evidence["criterion_high_kME"] = evidence["kME"].abs() >= 0.80
    evidence["criterion_robust_annotation"] = evidence["annotation_confidence"].isin(
        ["moderate-to-high homology-supported", "moderate homology-supported"])
    criterion_cols = [c for c in evidence.columns if c.startswith("criterion_")]
    evidence["criteria_met_n"] = evidence[criterion_cols].sum(axis=1)
    evidence["integrated_evidence_class"] = np.where(evidence[criterion_cols].all(axis=1), "Tier A",
                                                       np.where(evidence["criteria_met_n"] >= 3, "Tier B", "Tier C"))
    write_tsv(evidence, TABLES / "11_Genome_to_physiology_starch_gene_evidence.tsv")

    # Figures.
    plot_transition(transition_summary)
    plot_enrichment(nonredundant)
    plot_wgcna(wgcna)
    plot_conservation(conservation)

    # Input/QC manifests.
    manifest_rows = []
    for path in sorted(required):
        manifest_rows.append({"path": str(path.relative_to(ROOT)), "bytes": path.stat().st_size,
                              "sha256": sha256(path)})
    input_manifest = pd.DataFrame(manifest_rows)
    write_tsv(input_manifest, QC / "INPUT_MANIFEST_SHA256.tsv")

    max_kme_delta = float(np.nanmax(np.abs(all_kme["kME"] - all_kme["retained_own_module_kME"])))
    selected_kme = all_kme[all_kme["gene_id"].isin(selected_ids)].copy()
    selected_max_kme_delta = float(np.nanmax(
        np.abs(selected_kme["kME"] - selected_kme["retained_own_module_kME"])))
    qc_summary = {
        "status": "PASS_WITH_DECLARED_LIMITATIONS",
        "developmental_analysis_source": "fresh final-v3-full DESeq2 and WGCNA recomputation",
        "developmental_concordance_status": concordance_audit["status"],
        "vst_recompute_max_absolute_difference": vst_max_abs_delta,
        "vst_recompute_flattened_Pearson_r": vst_flat_correlation,
        "lrt_recompute_BH05_classification_agreement": lrt_bh05_agreement,
        "final_v3_gene_count_in_vst": int(len(dev_vst)),
        "developmental_samples": len(dev_samples),
        "tissue_samples": len(tis_samples),
        "figure4_gene_count": len(selected),
        "author_id_mapping_count": len(mapped),
        "unmapped_new_v3_models": int(len(id_map) - len(mapped)),
        "go_bp_terms_mapped": len(go_terms),
        "kegg_pathways_mapped": len(kegg_terms),
        "go_annotated_final_v3_genes": len(set().union(*go_terms.values())),
        "kegg_annotated_final_v3_genes": len(set().union(*kegg_terms.values())),
        "wgcna_assigned_genes": len(assignments),
        "wgcna_kme_genes_recomputed": len(all_kme),
        "wgcna_assignment_genes_absent_from_current_vst": len(assignment_not_in_vst),
        "current_vst_genes_without_wgcna_assignment": len(vst_without_assignment),
        "current_vst_genes_without_assignment_zero_mad": vst_without_assignment_zero_mad,
        "current_vst_genes_without_assignment_positive_mad": len(vst_without_assignment) - vst_without_assignment_zero_mad,
        "assigned_genes_with_zero_mad_in_current_vst": assigned_in_vst_zero_mad,
        "max_abs_recomputed_vs_retained_kME_difference": max_kme_delta,
        "selected_gene_max_abs_Pearson_vs_retained_bicor_kME_difference": selected_max_kme_delta,
        "selected_genes_with_unique_orthogroup": int(conservation["orthogroup"].notna().sum()),
        "selected_combined_1to1_syntenic_orthologs": int((conservation["combined_status"] == "1:1 syntenic ortholog").sum()),
        "selected_combined_syntenic_duplicated_orthologs": int((conservation["combined_status"] == "syntenic duplicated ortholog").sum()),
        "protein_identity_coverage_status": "not computed; retained release lacks pairwise alignment identity/coverage fields",
        "local_sv_status": "skipped; no retained high-confidence event-level SV dataset covering all 20 loci",
    }
    (QC / "QC_SUMMARY.json").write_text(json.dumps(qc_summary, indent=2) + "\n")

    strongest = transition_summary.sort_values("n_total_DEG", ascending=False).iloc[0]
    go_sig = int((go_enrichment["q_value"] < FDR).sum())
    kegg_sig = int((kegg_enrichment["q_value"] < FDR).sum())
    lxy_status = Counter(conservation["LXY_status"])
    bun_status = Counter(conservation["BunLong_status"])
    combined_status = Counter(conservation["combined_status"])
    report = f"""# GY2 genome-to-physiology final incremental analysis report

## 1. Input inventory

- Final GY2 v3 developmental expression: PRJNA756190, 18 samples (6 stages x 3 replicates).
- Final GY2 v3 tissue expression: PRJNA1493089, 12 samples (4 tissues x 3 replicates).
- Fresh final-v3-full developmental DESeq2 contrasts/LRT/VST recomputed from the frozen 20,063-gene count matrix; read mapping and count summarization were not rerun.
- Fresh WGCNA assignments/eigengenes/stage summary rebuilt from the matching 18-sample final-v3-full VST matrix.
- Fixed Figure 4 pathway set: 20 final `GY2U` genes.
- Final-v3 annotation coordinates/proteins, seven-species Orthogroups, and GY2-LXY/GY2-Bun Long MCScanX anchors.
- Vendor functional annotation transferred only through the exact one-to-one author-ID map.
- The earlier v3-primary contrast/WGCNA pipeline was superseded and was not used in final statistics.

All input paths, sizes, and SHA-256 values are in `qc/INPUT_MANIFEST_SHA256.tsv`.
Fresh final-v3-full VST values matched the independently retained task109 matrix (maximum absolute difference {vst_max_abs_delta:.3g}; flattened Pearson r = {vst_flat_correlation:.12g}). LRT q < 0.05 classification agreement was {lrt_bh05_agreement:.1%}; detailed numeric deltas are in `qc/FINAL_V3_FULL_RECOMPUTE_CONCORDANCE.json`.

## 2. Analysis versions

- Python {platform.python_version()}
- pandas {pd.__version__}
- numpy {np.__version__}
- scipy imported from the execution environment
- matplotlib {matplotlib.__version__}
- R 4.5.1; DESeq2 1.50.2; WGCNA 1.74 (server recomputation; see `sessionInfo.txt`).
- WGCNA soft power 16 (scale-free R2 = 0.808762; mean connectivity = 54.4691), selected by the preregistered smallest-R2>=0.80 and mean-connectivity>=5 rule.
- Thresholds: BH q < {FDR}; absolute log2FC >= {LFC} for adjacent DEGs.

## 3. Developmental transition results

The strongest adjacent transition was {strongest['transition']} with {int(strongest['n_total_DEG']):,} DEGs ({int(strongest['n_up']):,} up, {int(strongest['n_down']):,} down). Full counts are in `01_Adjacent_stage_DEG_transition_summary.tsv`.
The DESeq2 inclusion universe contained 17,174 genes; 17,142 had finite p values in every adjacent-stage contrast, while the same 32 genes had NA inferential values and were never called differential.

## 4. Transition functional enrichment

GO Biological Process and KEGG enrichment used Fisher exact tests against genes that were both testable in each corresponding contrast and represented in the respective transferred GO/KEGG annotation, followed by BH correction within each transition/direction/database analysis. Terms required at least five background genes and two foreground hits. There were {go_sig:,} significant GO rows and {kegg_sig:,} significant KEGG rows at q < 0.05. GO redundancy was reduced by retaining the lowest-q representative when foreground-gene Jaccard overlap was >= 0.70.

## 5. Corm-enriched genes

- Strict corm-enriched: {int(tissue_all['strict_corm_enriched'].sum()):,}
- Additional relaxed corm-biased: {int(tissue_all['relaxed_corm_biased'].sum()):,}

All reported effects are corm-vs-other-tissue log2 fold changes; the retained final-v3 tissue contrast files were stored as other-tissue-vs-corm and were sign-inverted explicitly.

## 6. Corm-developmental intersection

- Development-responsive (LRT q < 0.05): {int(dev_lrt['development_responsive'].sum()):,}
- Strict corm-enriched: {int(tissue_all['strict_corm_enriched'].sum()):,}
- Intersection: {len(corm_dev):,}
- Figure 4 pathway overlap: {a} of {len(pathway_set)} genes; Fisher OR = {pathway_or:.4g}, p = {pathway_p:.4g}.

This test is enrichment evidence only; a nonsignificant result is not reinterpreted as pathway confirmation.

## 7. WGCNA-pathway integration

kME was recomputed as Pearson correlation between each eligible gene's 18-sample VST vector and its assigned module eigengene after exact sample reconciliation. BH correction was applied across all {len(all_kme):,} recomputed eligible non-grey gene tests. {int(wgcna['kME'].notna().sum())} of 20 pathway genes had non-grey kME estimates; {int(wgcna['high_module_membership'].fillna(False).sum())} met the descriptive high-membership criterion (absolute kME >= 0.80 and q < 0.05). Module enrichment used all WGCNA-eligible genes as background; significance is reported without forcing a positive result.

The developmental VST and WGCNA now come from the same final-v3-full recomputation: {len(assignment_not_in_vst):,} assignment genes are absent from the VST, while {len(vst_without_assignment):,} VST genes lack an assignment ({vst_without_assignment_zero_mad:,} zero-MAD and {len(vst_without_assignment) - vst_without_assignment_zero_mad:,} positive-MAD). Thus the unassigned genes are the explicit zero-MAD exclusions from network construction. All 20 fixed pathway genes are present and uniquely assigned. The WGCNA own-module statistic used bicor, whereas the requested integration statistic here is Pearson; bicor is retained only as an audit column and is not claimed to be numerically identical.

## 8. Three-genome pathway conservation

- LXY status counts: {dict(lxy_status)}
- Bun Long status counts: {dict(bun_status)}
- Conservative combined status counts: {dict(combined_status)}

Statuses integrate final-v3 Orthogroups and retained MCScanX anchors. A strict 1:1 label additionally requires exactly one in-orthogroup anchor and no extra cross-orthogroup anchor; this conservatively classifies `GY2U013960` as syntenic duplicated in both comparisons. Copy numbers are orthogroup protein-model counts, not population CNV calls. Protein identity and coverage are left blank because pairwise alignment identity/coverage fields were not retained; MCScanX e-values were not repurposed as identity.

## 9. Local SV context

Not performed. The retained release does not contain a high-confidence, event-level SV dataset with complete locus-linked coverage for all 20 final-v3 genes. No read-backed SV, SV-expression causality, or absolute absence is claimed.

## 10. Integrated evidence matrix

`11_Genome_to_physiology_starch_gene_evidence.tsv` combines coordinates, orthogroups, comparator status, developmental LRT, developmental amplitude/peak, WGCNA module/kME, and three tissue contrasts. Tier A/B/C is a transparent count of five stated evidence criteria, not a causal score or validation ranking.

## 11. Main-text use

- Use the adjacent-transition summary as Figure 3D.
- State that transcriptional reprogramming intensifies across development only to the extent supported by the fresh final-v3-full recomputed contrasts.
- Describe combined developmental/corm-biased pathway genes as associated candidates.
- Describe orthogroup/MCScanX results as comparative genome context, not cultivar-causal evidence.

## 12. Supplementary use

- Transition-specific GO/KEGG enrichment.
- Pathway-gene WGCNA membership and module enrichment.
- Three-genome orthogroup/copy/synteny status.
- Functional annotation QC and all-gene corm classifications.

## 13. Explicit limitations

1. Functional annotations for 18,990 inherited models were transferred through an exact one-to-one final-v3/author-ID map; {int(len(id_map) - len(mapped)):,} new/changed v3 models lacked an author-ID mapping and therefore were not assigned inherited GO/KEGG terms.
2. Swiss-Prot-like symbols are homology-based annotations, not experimental functional confirmation in taro.
3. Orthogroup copy counts are annotation/protein-model counts and are sensitive to model fragmentation.
4. Pairwise protein identity/coverage was not retained and was not inferred from e-values.
5. No complete read-backed local SV dataset was available for the 20 loci.
6. No experimental qPCR validation dataset was available; none was simulated.
7. All expression, WGCNA, tissue-bias, and comparative associations are non-causal.
8. WGCNA excluded zero-MAD genes before network construction; the exact exclusion count is quantified in `qc/QC_SUMMARY.json` and none of the 20 fixed pathway genes was removed.
"""
    (ROOT / "GY2_genome_to_physiology_FINAL_ANALYSIS_REPORT.md").write_text(report)

    readme = """GY2 genome-to-physiology final incremental analysis

Run:
  python3 scripts/integrated_expression_analysis.py

This package uses final GY2 v3 processed expression, a fresh final-v3-full
developmental DESeq2/WGCNA recomputation, final-v3 Orthogroups, and MCScanX
anchors. It does not rerun assembly, annotation, STAR, featureCounts,
whole-genome alignment, or SV calling.

The earlier v3-primary developmental contrast/WGCNA pipeline is superseded and
is intentionally excluded from the final release package.

Results:
  results/tables/   final tables
  results/figures/  600-dpi LZW TIFF figures and PNG previews
  qc/               input checksums and QC summary
  scripts/          reproducible analysis script

Interpretation boundaries:
  No causal gene claims, no read-backed SV claims, no absolute gene-absence
  claims, no population pangenome claims, and no simulated qPCR.
"""
    (ROOT / "README_FINAL_ANALYSIS.txt").write_text(readme)

    output_rows = []
    output_manifest_path = QC / "OUTPUT_MANIFEST_SHA256.tsv"
    for path in sorted([p for p in ROOT.rglob("*") if p.is_file() and ".extract" not in p.parts]):
        if path.is_relative_to(INPUT):
            continue
        if path == output_manifest_path:
            continue
        output_rows.append({"path": str(path.relative_to(ROOT)), "bytes": path.stat().st_size,
                            "sha256": sha256(path)})
    write_tsv(pd.DataFrame(output_rows), output_manifest_path)

    print(json.dumps({
        "status": "complete_with_declared_limitations",
        "tables": len(list(TABLES.glob("*.tsv"))),
        "tiff_figures": len(list(FIGURES.glob("*.tiff"))),
        "strongest_transition": strongest["transition"],
        "strict_corm_enriched": int(tissue_all["strict_corm_enriched"].sum()),
        "corm_developmental": len(corm_dev),
    }, indent=2))


if __name__ == "__main__":
    main()
```
