## Supplementary material for "A chromosome-scale genome and transcriptomic atlas of taro (Colocasia esculenta) corm development": Data S2: README.md

# Data S2. Figure source data

| Figure | Data files |
| --- | --- |
| Figure 1 | Chromosome lengths, genomic tracks, syntenic anchors and repeat composition; Hi-C contact map |
| Figure 2 | (A) OrthoFinder species tree, (B) gene categories, (C) chromosome correspondences, (D) NG86 Ks distributions and (E) CAFE5 family changes |
| Figure 3 | (A, B) Developmental and tissue PCA source values, (C) WGCNA module summaries and (D) adjacent-stage DEG counts |
| Figure 4 | The 20-gene carbohydrate panel, pathway annotations, developmental and tissue expression values, and cross-genome conservation |

Tab-delimited files are UTF-8 encoded. Gene identifiers correspond to the GY2 v3 annotation in Data S1.
