## Supplementary figures and images for "A chromosome-scale genome and transcriptomic atlas of taro (Colocasia esculenta) corm development"

### HiC_contact_map.png

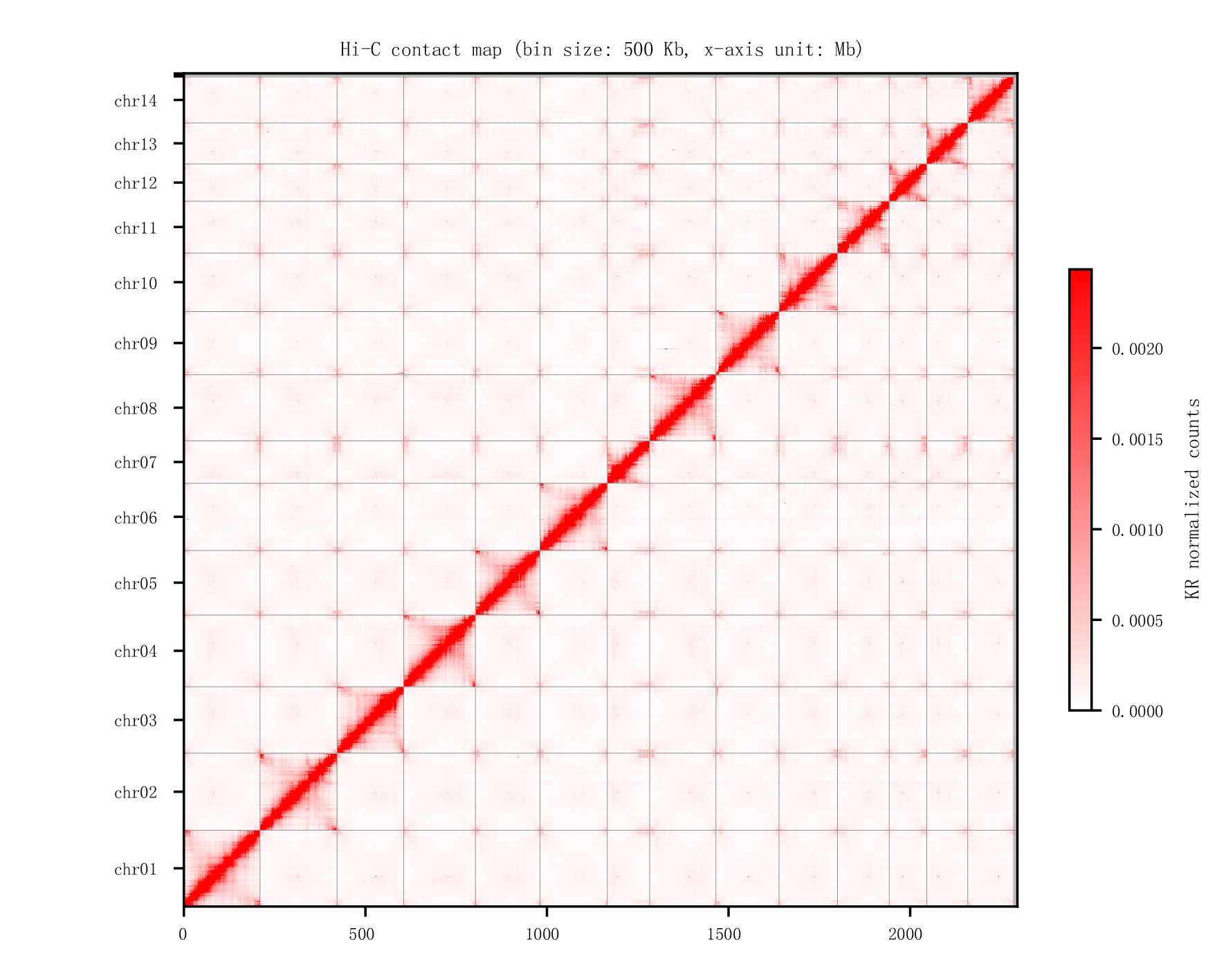
